# Gene expression-based identification of antigen-responsive CD8^+^ T cells on a single-cell level

**DOI:** 10.1101/673707

**Authors:** Yannick F. Fuchs, Virag Sharma, Anne Eugster, Gloria Kraus, Robert Morgenstern, Andreas Dahl, Susanne Reinhardt, Andreas Petzold, Annett Lindner, Doreen Löbel, Ezio Bonifacio

**Author notes:** these authors contributed equally to the work (Yannick F. Fuchs, Virag Sharma).

## Abstract

CD8^+^ T cells are important effectors of adaptive immunity against pathogens, tumors and self antigens. Here, we asked how human cognate antigen-responsive CD8^+^ T cells and their receptors could be identified in unselected single-cell gene expression data. Single-cell RNA sequencing and qPCR of dye-labelled antigen-specific cells identified large gene sets that were congruently up- or downregulated in virus-responsive CD8^+^ T cells under different antigen presentation conditions. Combined expression of *TNFRSF9*, *XCL1*, *XCL2*, and *CRTAM* was the most distinct marker of virus-responsive cells on a single-cell level. Using transcriptomic data, we developed a machine learning-based classifier that provides sensitive and specific detection of virus-responsive CD8^+^ T cells from unselected populations. Gene response profiles of CD8^+^ T cells specific for the autoantigen islet-specific glucose-6-phosphatase catalytic subunit-related protein differed markedly from virus-specific cells. These findings provide single-cell gene expression parameters for comprehensive identification of rare antigen-responsive cells and T cell receptors.

**One-sentence summary:** Identification of genes, gene sets, and development of a machine learning-based classifier that distinguishes antigen-responsive CD8^+^ T cells on a single-cell level.

## Introduction

CD8^+^ T cells are integral to the clearance of virus-infected cells and the control of cell transformation. These attributes are exploited by therapies, such as vaccination against infection and immune therapies targeting cancer. CD8^+^ T cells are also involved in the destruction of cells in some autoimmune diseases, such as type 1 diabetes, and in graft-versus-host disease. In active immune responses, CD8^+^ T cells undergo rapid clonal expansion and this expansion of activated CD8^+^ T cells can be a marker of ongoing infection in immune-mediated disease. Therefore, it is conceivable that the detection and information provided by clonal expansions can be used to identify target antigens or disease-causing agents, and to develop therapies that exploit CD8^+^ T cell specificity to control disease.

CD8^+^ T cell specificity is provided by T cell receptors (TCR), which recognize a cognate antigen peptide presented by major histocompatibility complex (MHC) class I molecules on the cell surface. This mechanism of antigen recognition has been used to develop fluorochrome-labeled, peptide-loaded MHC class I multimers and has led to sophisticated methods combining multiple fluorochromes and peptides to find and isolate antigen-specific CD8^+^ T cells (1–3). These reagents also allow detailed TCR and phenotypic analysis of cells using single-cell technologies (4–6). Multimers require high levels of TCR expression for the detection of antigen-specific cells (7). Although high TCR expression is a feature of CD8^+^ T cells in resting conditions, CD8^+^ T cells undergo numerous changes in gene and protein expression upon stimulation with their target peptide (8). In particular, downregulation of TCR is a consistent response to the engagement of the cognate peptide, conceivably for negative feedback control of T cell activity (9–11). Therefore, the selection of multimer-binding CD8^+^ T cells may bias our understanding of the phenotype of antigen-specific CD8^+^ T cells.

We have incorporated MHC class I multimers and peptide activation into an unbiased approach to analyze antigen-specific CD8^+^ T cells from peripheral blood. Here, we demonstrate marked downregulation of TCR upon stimulation with the cognate peptide and describe protein and gene sets that can be used to identify and isolate antigen-responsive CD8^+^ T cells. We show how these approaches can be combined with MHC multimers to obtain a comprehensive description of activated and non-activated antigen-specific CD8^+^ T cells, and how they can be used without MHC multimers for identification and in-depth gene expression profiling of cells responding to their cognate antigen. Additionally, we provide evidence that multimer binding (specificity) and peptide responsiveness are not identical.

## Results

### Identification of highly pure Flu MP_58-66_-responsive CD8^+^ T cells for single-cell analysis

The activation of influenza matrix protein 1_58-66_ (Flu MP_58-66_)-specific CD8^+^ T cells by presentation of their cognate peptide compromised the ability to detect these cells using MHC class I multimers (Fig. 1a), presumably due to downregulation of TCR (Fig. S1). Therefore, we suspected that isolation of multimer-positive CD8^+^ T cells may miss recently activated cells, and we examined whether it was possible to complement MHC class I multimers with activation markers. To identify potential activation markers on the gene expression level, antigen-specific CD8^+^ T cells were isolated from carboxyfluorescein succinimidyl ester (CFSE)-labeled peripheral blood mononuclear cells (PBMCs) via Flu MP_58-66_ human leukocyte antigen (HLA) class I A*0201 multimers and directly flow sorted to unlabeled antigen-presenting cells (K562/A*0201 cells or autologous PBMCs; schematic in Fig. 1b). Cells were stimulated overnight with the cognate peptide, mock peptide, or solvent, and CFSE-labeled cells were flow sorted as single cells (Fig. 1c) and the gene expression of 75 selected genes analyzed. Consistent with the impaired detection of antigen-responsive cells via multimers when cells had been exposed to their cognate peptide, the multimer fluorescence intensity of dye-labeled cells incubated with their cognate antigen was lower than that of cells incubated with solvent or mock peptide (Fig. 1c, top). Cells stimulated with the cognate peptide had distinct gene expression profiles compared with the control stimulated cells for each of three donors and either K562/A*0201 or PBMCs as the antigen-presenting cells (T-distributed stochastic neighbor embedding [t-SNE] plots, Fig. 1c). As expected, the genes encoding known activation markers (CD137 [*TNFRSF9*], CD69, and CD25) and effector molecules (*IFNG*, *GZMB*, and *FASLG*) were upregulated in cells incubated with their cognate peptide Flu MP_58-66_ (Fig. 1c; Fig. S2) compared with cells incubated with mock peptide or solvent. Among the marker genes that best discriminated the cognate peptide-activated cells, upregulation of *TNFRSF9*, *REL*, *EGR2*, *SRP14*, *FASLG*, *GZMB*, *IFNG*, *CD69*, *HMGB1*, and *NAB2* and downregulation of *IL7R* were consistently detected in each of the donors (Table S1). Notably, the expression of some genes, including *IFNG*, was extremely heterogeneous between the responsive cells. Only minor differences were observed when K562/A*0201 cells were used as antigen-presenting cells compared with PBMCs (Fig. S2), and the cognate peptide stimulated Flu MP_58-66_-specific CD8^+^ T cells clustered together in the t-SNE analysis of the gene expression regardless of the type of antigen-presenting cell (Fig. 1c). We also found that the profiles of a small number of cognate peptide-stimulated cells were similar to those of the mock peptide- and solvent-stimulated cells (e.g., donor #3), potentially reflecting heterogeneity among the pool of antigen-directed cells.

**Fig.1:**
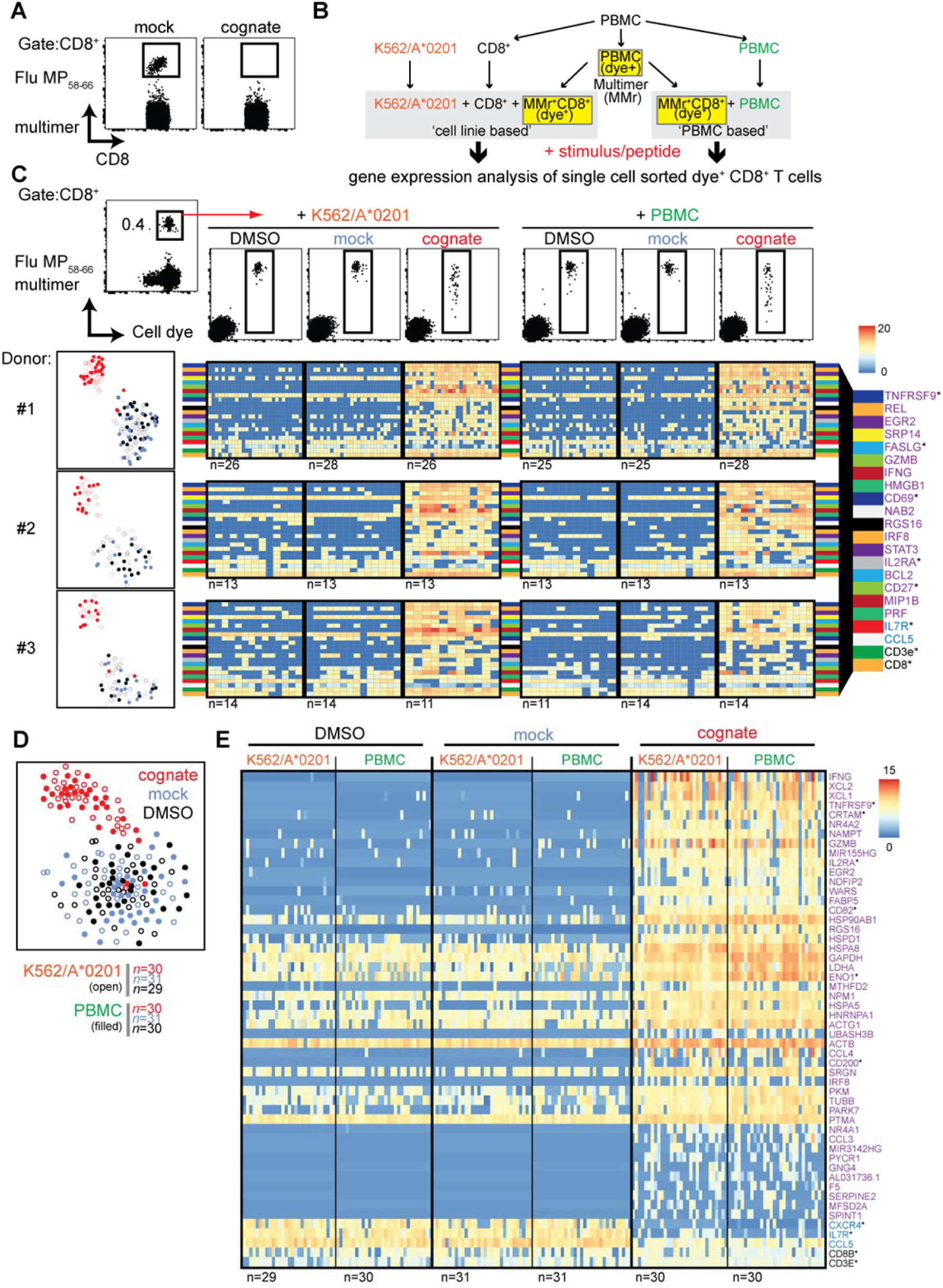
Single-cell gene expression analysis of sorted Flu MP_58-66_-responsive CD8^+^ T cells. (**A**) Representative flow cytometry dot plots of PBMCs stained with HLA-A2 multimers loaded with Flu MP_58-66_ after incubating PBMCs for 20 h in the presence of mock (IGRP_265-273_) or cognate (Flu MP_58-66_) peptide. Plots show 5 × 10^4^ cells in the CD8 gate. (**B**) Schematic work-flow of the dye-based activation assay. (**C**) *Top*: representative dot plots (donor #1) of multimer and cellular dye-stained PBMCs (left). Cells in the CD8 gate are shown. CD8 cells staining positive for the multimer and cell dye were sorted (red arrow) for use in assays using the K562/A*0201 cell line or autologous PBMCs for antigen presentation. After incubation with control stimuli (peptide solvent and mock peptide) or the cognate peptide for 20 h, CD8^+^ T cells staining positive for the cell dye were sorted for single-cell targeted gene expression analysis. *Lower left*: t-SNE analysis for donors #1–3. Gene expression was analyzed in single cells following incubation with an antigen-presenting cell line (open circles) or PBMCs (filled circles) in the presence of solvent (black) or mock peptide (blue) as control stimuli or with the cognate peptide (red). *Lower right*: Heatmaps of the top 20 ranked differentially expressed genes in cognate peptide-stimulated cells relative to control-stimulated cells for donors #1–3. The numbers of analyzed cells are shown below the individual heatmaps. Genes marked with an asterisk encode proteins expressed on the cell surface. (**D**) t-SNE of scRNAseq gene expression data for Flu MP_58-66_-directed CD8^+^ T cells derived from donor #1 and incubated with K562/A*0201 (open) or autologous PBMCs (filled) in the presence of solvent (black) or mock peptide (blue) as control stimuli or with cognate peptide (red). (**E**) Heatmaps for the top 50 ranked differentially expressed genes in antigen-directed CD8^+^ T cells incubated with cognate peptide relative to control stimuli (purple: upregulated genes; blue: downregulated genes; CD8B and CD3E are also shown). Data are shown for donor #1 following incubation with K562/A*0201 or autologous PBMCs. Genes were combined for the ranking. The numbers of analyzed cells are shown below the heatmaps.

The findings using the target qPCR expression were extended using single-cell RNA sequencing (scRNAseq) of cells from donor #1 that were processed and isolated in the same manner. The TCR sequences of the Flu MP_58-66_ multimer-positive cells were dominated by *TRAJ42*01* (66.5%) and *TRBV19*01* (95.7%), as previously described for Flu MP_58-66_-directed TCRs (4, 12–14) (Table S2). Again, the cognate peptide-stimulated cells were distinct from the mock peptide- and solvent-stimulated cells, and only a minority of cognate peptide-stimulated cells had gene expression profiles that were not distinguishable from the mock peptide- and solvent-stimulated cells (Fig. 1d). The TCR sequences of these non-responsive cells included TCRs that were also detected in responsive cells, which suggests that these were unresponsive Flu-specific CD8^+^ T cells rather than an isolation artefact. In total, 2360 genes were differentially expressed (adjusted *p* < 0.05) upon stimulation with the cognate peptide relative to mock peptide or solvent stimulation (Fig. S3; Table S3), of which 1940 had a >2 log_2_-fold change. Of these, 590 genes were differentially expressed (adjusted *p* < 0.05) in both K562/A*0201 cell- and PBMC-based peptide presentation. The top 50 differentially expressed genes are shown in Fig. 1e. They include *TNFRSF9*, *CCL5*, *CCL4*, *EGR2*, *GZMB*, *IFNG*, *IL2RA*, and *IL7R*, which were also observed using the targeted gene panel. They also comprised genes encoding the cytokines X-C motif chemokine ligand (XCL)1 and XCL2 or cytotoxic and regulatory T cell molecule (CRTAM) that are involved in attraction (15) and adhesion to cross-presenting dendritic cells (16), genes important for metabolic function (e.g., *GAPDH*, *FABP5*, (17)), and several other genes involved in protein synthesis supporting cell activation and expansion.

### Verification of the identified marker genes in CMVpp65_495-503_-responsive CD8^+^ T cells

To validate our findings obtained using Flu MP_58-66_-specific CD8^+^ T cells, we performed similar experiments using CD8^+^ T cells specific to the dominant human cytomegalovirus (hCMV) structural protein pp65 (CMV pp65_495-503_). CFSE-labeled multimer-isolated CMV-specific CD8^+^ T cells were incubated with PBMCs loaded with CMV pp65_495-503_ peptide or control antigen, and the CFSE-labeled cells were subsequently sorted for scRNAseq. The TCR repertoire of the isolated single cells resembled that expected for CMVpp65_495-503_-specific CD8^+^ T cells, and included the previously described enriched combinations of *TRAJ49*/*TRAV24* (donors #4–#6), *TRAJ50*/*TRAV35* (donors #4 and #5), *TRBV6-5*/*TRBJ1-2* (donors #4 and #5), and *TRBV12-4*/*TRBJ1-2* (donors #4 and #6) ((5); Table S2). Again, cells stimulated with the cognate peptide were separated from the control-stimulated cells based on their gene expression profiles (Fig. 2a), and the genes *TNFRSF9* (*CD137*), *XCL1*, *CRTAM*, *EGR2*, and *XCL2* were ranked highly as differentially expressed genes in all three donors (Fig. 2b). The expression of 2067 genes was increased (*n* = 1471) or decreased (*n* = 596) in cells stimulated with the cognate peptide in all three donors (Fig. S4; Table S4; Fig. 3a).

**Fig. 2:**
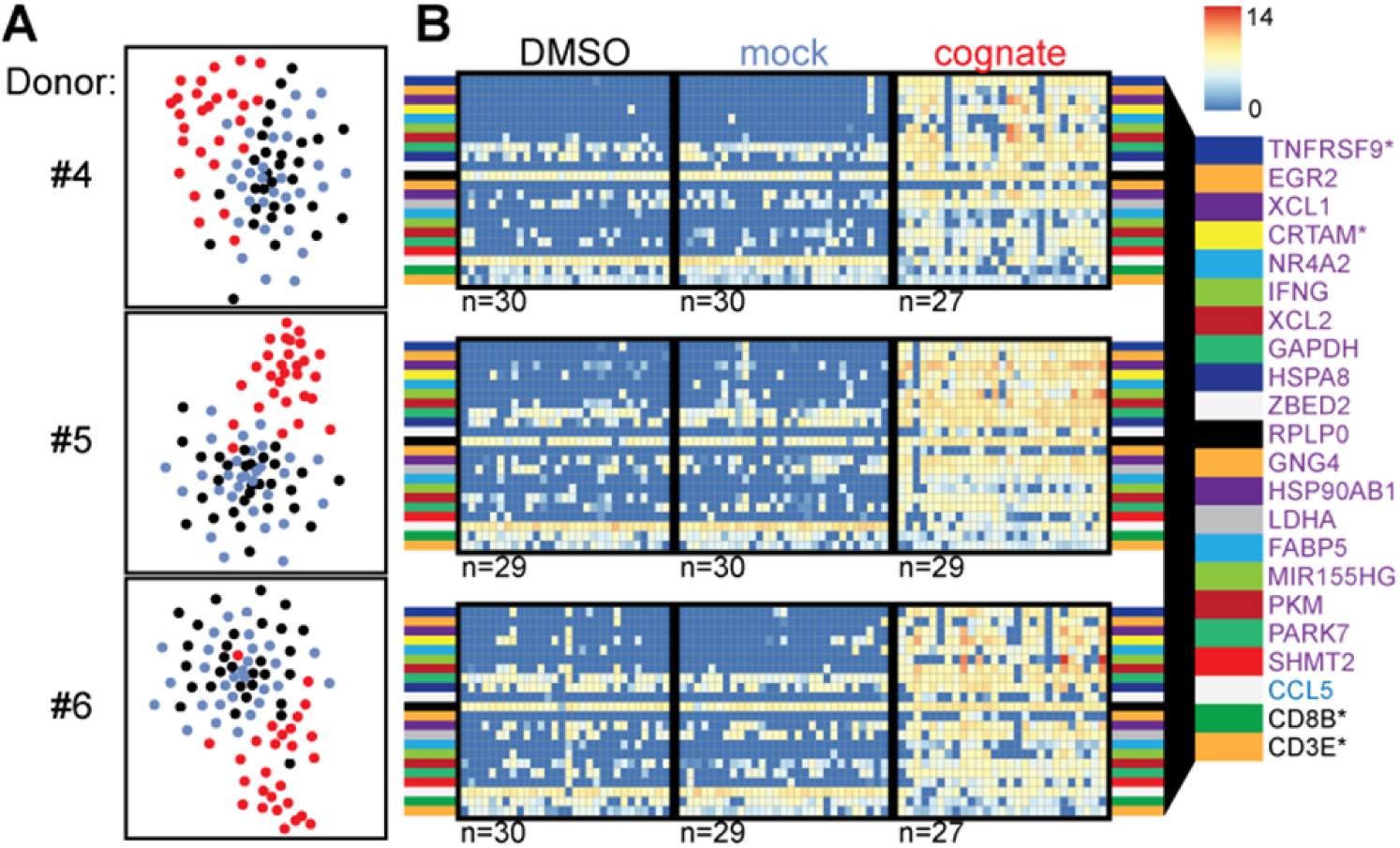
Verification of marker genes in CMVpp65_495-503_-responsive CD8^+^ T cells. (**A**) t-SNE analysis of scRNAseq data for CMVpp65_495-503_-directed CD8^+^ T cells from donors #4–6 following incubation with autologous PBMCs in the presence of solvent (black) or mock peptide (blue) as control stimuli or with the cognate peptide (red). (**B**) Heatmaps of the top 20 ranked differentially expressed genes in CMV pp65_495-503_-specific CD8^+^ T cells incubated with cognate peptide relative to control stimuli for donors #4–6 (purple: upregulated genes; blue: downregulated genes; CD8B and CD3E are also shown). The numbers of analyzed cells are shown below the heatmaps. Genes marked with an asterisk encode proteins expressed on the cell surface.

**Fig. 3:**
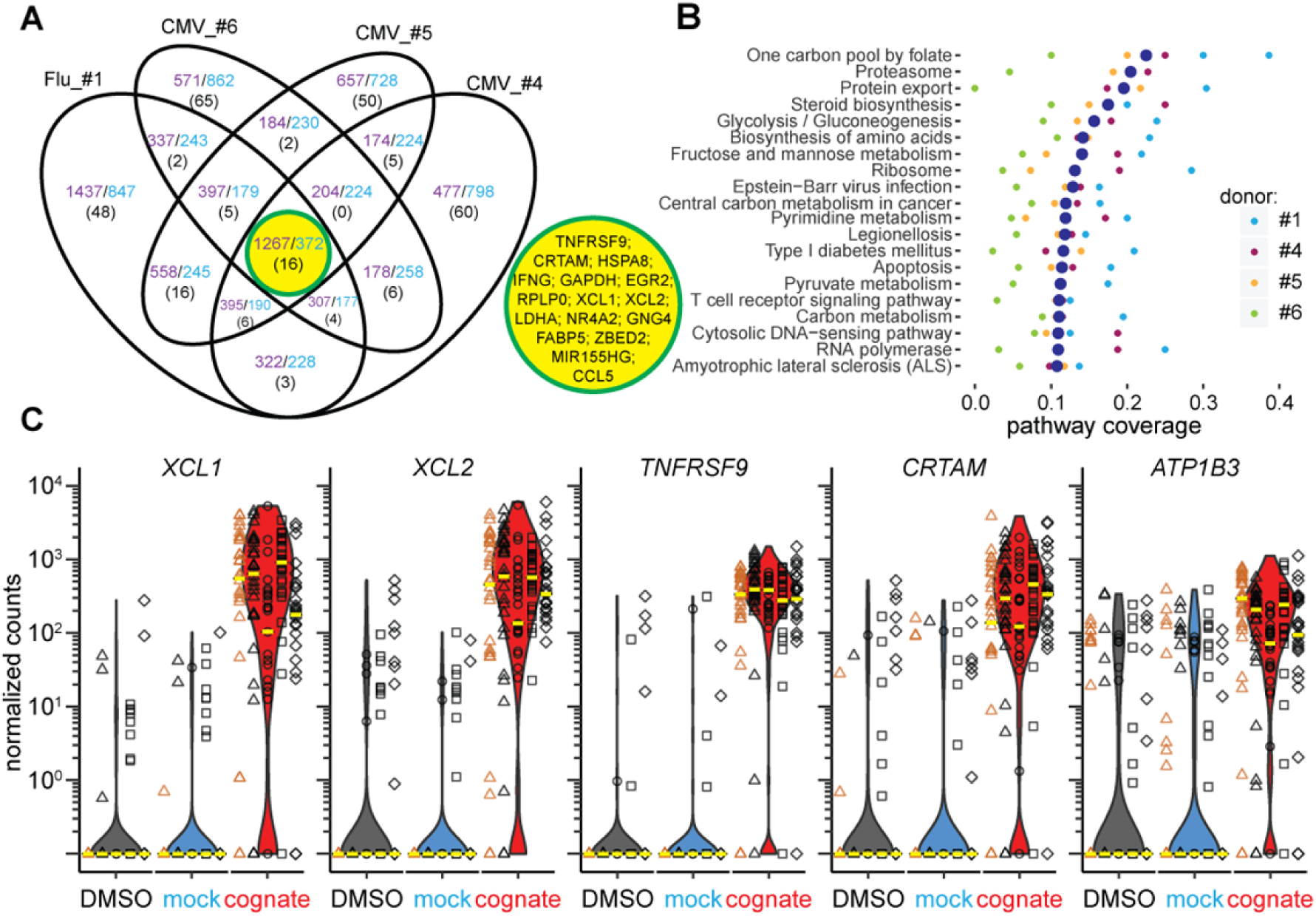
Combined gene and pathway analysis of Flu MP_58-66_- and CMVpp65_495-503_-responsive CD8^+^ T cells. (**A**) Venn diagram of overlapping genes upregulated (purple) or downregulated (blue) in Flu MP_58-66_-directed (donor #1) or CMVpp65_495-503_-directed CD8^+^ T cells (donors #4–6) following stimulation with the cognate peptide relative to control stimuli. The number of genes belonging to the top 100 differentially expressed genes in each individual is shown in parenthesis. Sixteen genes were shared between all individuals, as shown in the central yellow circle. (**B**) Top 20 KEGG pathways identified based on the differentially expressed genes in individual donors following stimulation with cognate peptide relative to control stimuli. The pathways showing significant enrichment for these genes in at least one donor were included in the analysis and were ranked based on the median pathway coverage (large blue dots) for all donors. (**C**) Scatter/violin plots showing normalized counts of the five genes that best separated the cognate peptide-stimulated and control-stimulated cells in antigen-directed CD8^+^ T cells incubated with cognate peptide (red violins) relative to solvent (black) or mock peptide (blue) in the presence of autologous PBMCs (black symbols) or K562/A*0201 (orange symbols). The *y*-axis represents the normalized read counts for individual Flu MP_58-66_-specific cells from donor #1 (triangles) or CMVpp65_495-503_-specific cells from donors #4 (circles), #5 (squares), and #6 (diamonds). The median values for individual donors and conditions are shown as yellow lines. Similar cell numbers were analyzed per condition for each donor, and data comprise counts for 148 cells incubated with DMSO, 152 cells incubated with mock peptide, and 143 cells stimulated with the cognate peptide.

### Identification of candidate marker genes common to Flu MP_58-66_-responsive and CMVpp_65495-503_-responsive CD8+ T cells

A total of 1639 genes were up- or downregulated in both the Flu MP_58-66_ -responsive (donor #1) and CMVpp65_495-503_-responsive (donors #4, #5, and #6) CD8^+^ T cells (Fig. 3a). We examined the top 100 differentially expressed genes in each of the donors and identified 16 that were shared between the CMV- and Flu-responsive cells. These included *TNFRSF9*, *CRTAM*, *XCL1*, *XCL2*, the transcription factors *EGR2* and *ZBED2,* the transcriptional regulator *NR4A2*, genes involved in metabolism (*GAPDH*, *LDHA*, and *FABP5*), the host gene for microRNA155 (*MIR155HG*), *RPLP0*, *HSPA8*, *GNG4*, *CCL5* and *IFNG*.

Pathway analysis of the differentially expressed genes in antigen-responsive cells across donors and antigen specificities repeatedly demonstrated the enrichment of genes associated with cell metabolism, infection, amino acid biosynthesis, apoptosis, TCR and cytokine signaling, protein processing, and antigen presentation (Fig. 3b).

We reasoned that markers that may be suitable for identifying activated antigen-responsive CD8^+^ T cells would be exclusively or most differentially expressed in either the cognate peptide-stimulated cells or the control mock peptide- and solvent-stimulated cells. Therefore, we ranked the genes according to their ability to separate these two groups (Table S5). The top 10 ranked genes were *TNFRSF9*, *FABP5*, *CRTAM*, *EGR2*, *NR4A2*, *XCL1*, *XCL2*, *ATP1B3*, *RBPJ* and *MIR155HG*. Among the top 50 separating genes, *XCL1*, *XCL2*, *TNFRSF9*, *CRTAM*, and *ATP1B3* showed the greatest differences in median expression levels between the cognate peptide- and control-stimulated cells (Fig. 3c). The ability to separate the two groups and the large differences in expression support the use of these genes as potential transcriptional markers for antigen-responsive CD8^+^ T cells.

We next assessed whether the top-ranking genes for separating the groups are also suitable for a wide range of peptide concentrations used for stimulation (Fig. S5). Even with a 1000-fold reduction in the peptide concentration, the expression of these genes differed markedly between mock-stimulated cells and cognate peptide-stimulated cells, suggesting that they mark antigen-responsive cells over a broad range of peptide concentrations.

### CD137, CD82 and CD355 surface proteins identify antigen-responsive cells

Several of the top-ranked differentially expressed genes encode cell-surface proteins. One of these, CD137 (encoded by *TNFRSF9*), is a known cell-surface marker of activated CD8^+^ T cells (18, 19). In accordance with our observations using Flu MP_58-66_-specific cells, the detection of CMV pp65_495-503_-directed cells via MHC class I multimers was impaired when the cells had been incubated with their cognate antigen. However, there was an increase in the frequency of CD137-stained CD8^+^ T cells (Figs. 4a; Fig. S6). We compared the TCR repertoires between CMV multimer-sorted CD8^+^ T cells from PBMCs stimulated with the control peptide (denoted antigen-specific CD8^+^ T cells) and CD137-positive CD8^+^ T cells from PBMCs stimulated with CMV pp65_495-503_ (denoted antigen-responsive cells). There was extensive overlap between the two repertoires (Fig. 4b; Table S2) indicating that both groups of cells display the same TCR specificity. As expected, there were marked differences in the gene expression profiles between antigen-specific and antigen-responsive cells, which included the expression of the key marker genes described above (e.g., *TNFRSF9*, *XCL1*, *XCL2*, and *CRTAM*; Fig. 4c). The differentially expressed genes broadly overlapped with those identified in cells stimulated with cognate antigen in the dye-based assay (Fig. 4d; Table S6), suggesting that multimer binding did not exert a major influence on the gene expression profiles in our dye-based CD8^+^ T cell activation assays. Additionally, the gene expression profiles of multimer-sorted antigen-specific cells resembled those of multimer-sorted CFSE-labeled CD8^+^ T cells incubated with control stimuli in previous assays (Fig. 4d).

**Fig. 4:**
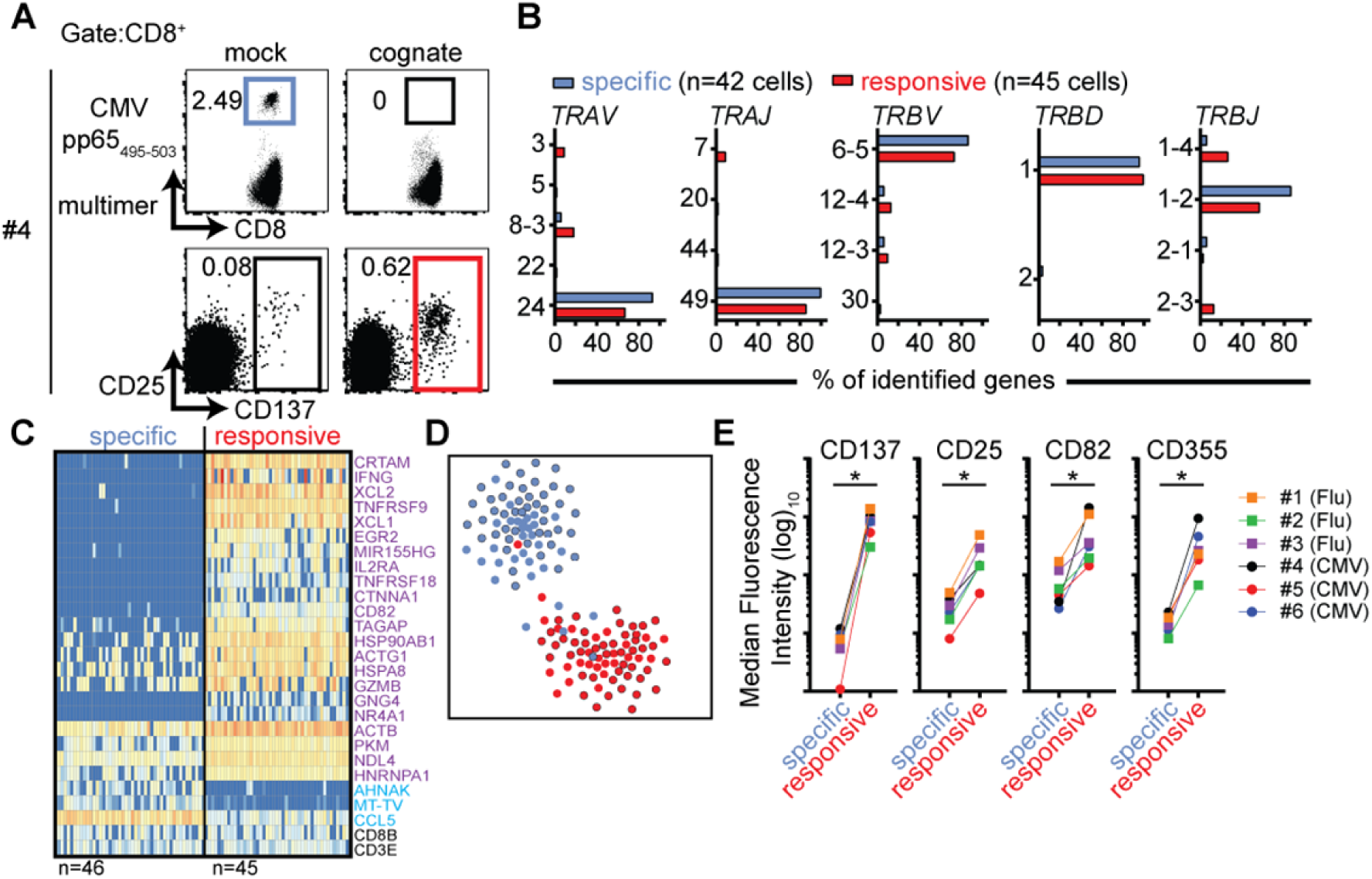
CD137 expression marks antigen-responsive CD8^+^ T cells in the absence of multimer staining. (**A**) Representative flow cytometry dot plots of PBMCs from donor #4 stained with HLA-A2 multimers loaded with CMVpp65_495-503_ peptide after incubating PBMCs for 20 h with mock (Flu MP_58-66_) or cognate (CMVpp65_495-503_) peptides (top) and corresponding plots showing CD25 and CD137 expression (bottom). All plots show 5 × 10^4^ cells in the CD8 gate. (**B**) TCR repertoire analysis of CD8^+^ T cells subjected to single-cell sorting based on positive staining with CMVpp65_495-503_ peptide-loaded HLA-A2 multimers from PBMCs stimulated with mock peptide (blue gate in **A**; antigen-specific) or based on CD137 expression after incubation with cognate peptide (representing cells in the red gate in **A**; antigen-responsive). The frequencies of genes used for the production of TCR α and β chains in each of the two populations of cells are shown. (**C**) Heatmaps of the top 25 ranked differentially expressed genes in CMV pp65_495-503_-specific relative to CMV pp65_495-503_-responsive CD8^+^ T cells from donor #4. The numbers of analyzed single cells are shown beneath the heatmaps. (**D**) t-SNE plot of gene expression data for single-cell-sorted antigen-specific (black outline filled blue) and antigen-responsive CD8^+^ T cells (black outline filled red). Data for CMV-directed cells incubated with mock (blue) or cognate (red) peptide in the dye-based activation assay are included as a reference. (**E**) Scatter plots of flow cytometric analyses comparing the median fluorescence intensities for the indicated surface markers between antigen-specific and antigen-responsive CD8^+^ T cells for donors #1–3 (donors with Flu-directed cells) and #4–6 (donors with CMV-directed cells).

Other genes encoding cell-surface proteins were also found among the genes that were differentially expressed in cells responding to their cognate antigen as compared with control-stimulated cells. We used antibodies directed against nine of these potential surface markers (CD72, CD82, CD134, CD160, CD200, CD319, CD357, CD355/CRTAM, and SEMA4A) and compared their surface protein expression levels in multimer-positive antigen-specific (mock peptide-exposed) and CD137^+^ antigen-responsive CD8^+^ T cells (cognate peptide-stimulated). In addition to CD137 and CD25, the expression levels of both CD82 and CD355 were consistently increased in antigen-responsive cells (Fig. 4e) for all six donors and both antigen specificities tested.

### Marker-based identification of antigen-responsive CD8^+^ T cells within unselected cells

To validate the identified gene expression markers of antigen-responsive cells further, we bulk-sorted CD95^+^CD8^+^ memory T cells from donor #4 after stimulating the cells with control (Flu MP_58-66_) or CMV pp65_495-503_ peptide, and we performed single-cell gene expression analysis and paired TCRα/β sequencing for over 4000 cells in each condition. Using the TCR information retrieved from CMV-specific cells in previous scRNAseq experiments of donor #4 (Table S2), we were able to trace the CMV-directed cells within the bulk population. Among control peptide-stimulated cells, the CMV-directed cells were not easily distinguishable from other CD8^+^ T cells based on their gene expression profiles (Fig. 5a). However, when stimulated with their cognate peptide, the CMV TCR-bearing cells in the CMV stimulation conditions almost exclusively clustered together and were clearly separated from the thousands of other CD8^+^ T cells based on their gene expression profiles. This cluster of cells was characterized by upregulation of the previously identified genes, including *XCL1*, *XCL2*, *TNFRSF9*, and *CRTAM* (Fig. 5b).

**Fig. 5:**
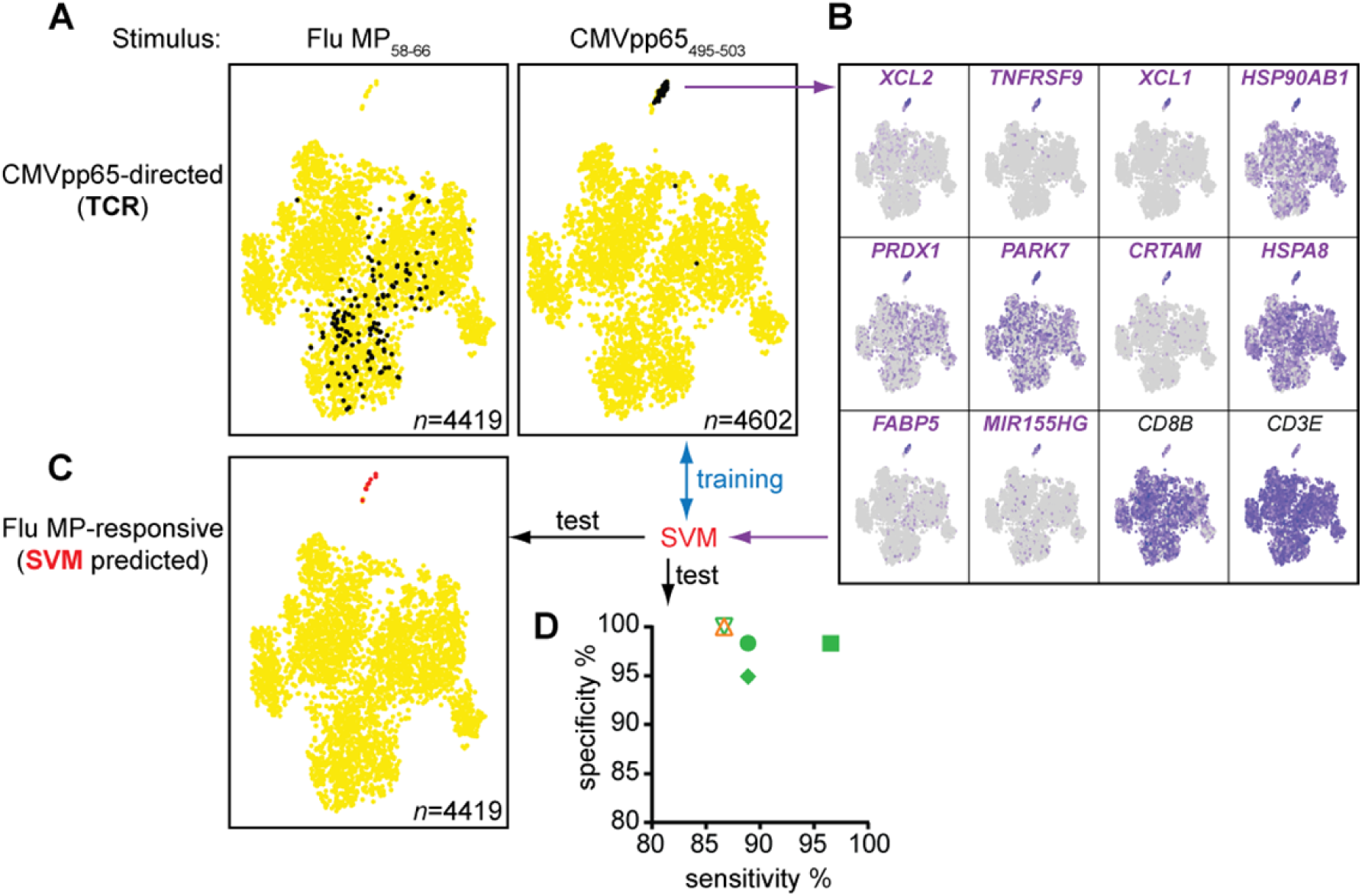
Multimer-independent identification of antigen-responsive cells from bulk memory CD8^+^ T cells. (**A**) t-SNE plots of gene expression data from CD95^+^CD8^+^ memory cells sorted from the PBMCs of donor #4 incubated for 20 h with the indicated peptides. Cells expressing CMVpp65_495-503_-directed TCRs, known from previous CMV multimer-based experiments, are shown in black. (**B**) t-SNE plots for CMV peptide-stimulated cells in (**A**) showing the expression levels of the top 10 genes marking the cluster of antigen-responsive cells and of the reference genes *CD3E* and *CD8B*. The 10 genes were used to train a machine learning algorithm (SVM), which was subsequently tested for its sensitivity and specificity to distinguish antigen-responsive cells from control stimulated-cells in the scRNAseq data sets of donors #4–6 (CMV-directed) and # 1 (Flu-directed) (**D**) and to predict Flu MP_58-66_-responsive cells from bulk memory CD95^+^CD8^+^ T cells (**C**).

Next, we used a machine learning algorithm (support-vector machine, SVM) to distinguish antigen-responsive from other cells using the top 10 marker genes identified in the cluster of antigen-responsive cells. The algorithm was trained using data obtained with the CMV peptide-stimulated CD95^+^CD8^+^ memory T cells to identify a hyperplane between antigen-responsive and non-responsive cells. We applied the algorithm to the scRNAseq datasets for donors #1–4. The algorithm correctly distinguished the cognate peptide-stimulated cells from the control-stimulated cells with high sensitivity (mean sensitivity 89.5%; range 86.7–96.6%) and specificity (mean 98.3%; range 94.9–100%) (Fig. 5d). We also tested the algorithm on the Flu MP_58-66_ peptide-stimulated CD95^+^CD8^+^ memory T cells from donor #4. Using a total of 4419 cells, the algorithm classified eight as antigen-responsive (Fig. 5c). All eight cells were in close proximity within the t-SNE analysis and where the cluster of responsive cells was expected. Paired TCR information was available for five of the eight cells in this dataset and included the previously described public Flu MP_58-66_-restricted TCR sequences (Table S7), suggesting that the algorithm can be used to identify the relevant antigen-specific and antigen-responsive CD8^+^ T cells.

Using this independent approach, we confirmed that the identified genes are potent marker genes. Additionally, we showed that it was possible to identify antigen-responsive cells within a bulk of memory CD8^+^ T cells, independently of MHC class I multimers.

### Autoantigen-specific CD8^+^ T cells may differ in their responsiveness

In addition to CD8^+^ T cells specific to viral antigens, we extended our study to autoreactive CD8^+^ T cells directed against the type 1 diabetes autoantigen islet-specific glucose-6-phosphatase catalytic subunit-related protein (IGRP). CD8^+^ T cells specific for the HLA-A*0201-restricted IGRP_265-273_ peptide were previously discussed as potential effectors of the islet destructive process in patients with type 1 diabetes and in HLA transgenic mice (20–22). Furthermore, IGRP_265-273_-directed CD8^+^ T cell clones were shown to kill peptide-loaded target cells in an antigen-specific manner (22, 23). We applied our dye-based CD8^+^ T cell activation assay to IGRP_265-273_-directed CD8^+^ T cells from a previously described patient with high frequencies of both Flu MP_58-66_- and IGRP_265-273_-directed cells (Fig. 6a) (23). In contrast to our findings with Flu- and CMV-directed cells, we did not observe a reduction in multimer fluorescence intensity or upregulation of CD137 protein after stimulation with cognate IGRP_265-273_ peptide in the dye-positive or dye-negative cells (Fig. 6b). As expected, CD137 was observed on the dye-negative CD8^+^ T cells stimulated with Flu MP_58-66_ peptide (mock), suggesting that the response to the viral peptide was not impaired in this patient. The dye-positive cells displayed a restricted TCR repertoire and the dominant IGRP TCR clonotype previously described for this donor (Table S2) (23). IGRP cells stimulated with cognate peptide did not cluster separately from control-stimulated cells (Fig. 6c), and only 321 genes were differentially expressed between cognate peptide- and control-stimulated cells, of which only six were shared between both types of antigen presentation. None of the top 50 differentially expressed genes were found with K562/A*0201- and PBMC-based antigen stimulation (Table S8; Fig. S7). None of the top 16 genes previously identified for virus-responsive CD8^+^ T cells were differentially expressed between the IGRP cognate peptide- and control-stimulated cells (Fig. 6d).

**Fig. 6:**
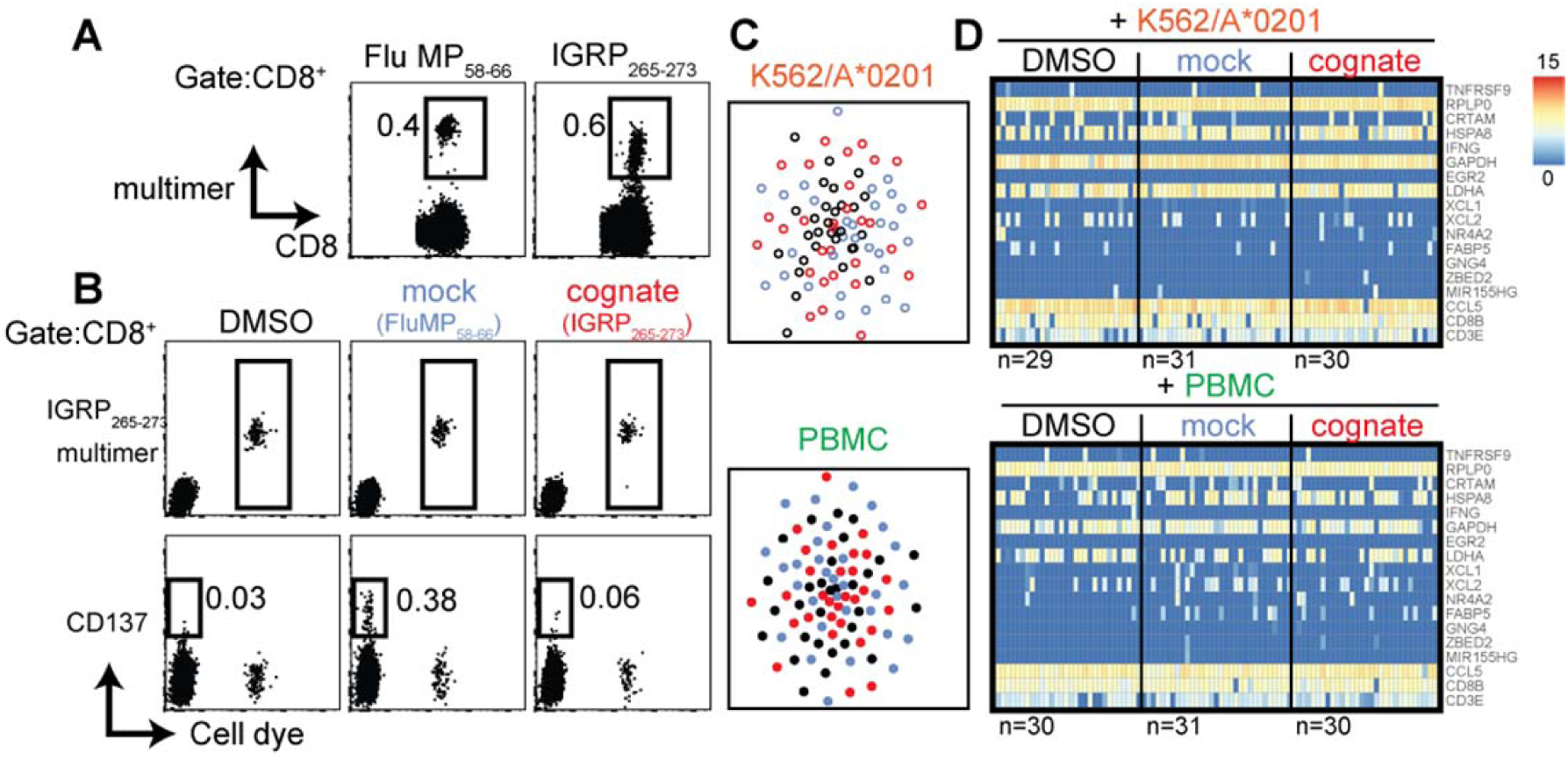
Autoantigen-directed CD8^+^ T cells may differ in their responsiveness to the cognate peptide. (**A**) Representative flow cytometry dot plots of PBMCs from a donor with type 1 diabetes stained with HLA-A2 multimers loaded with Flu MP_58-66_ or IGRP_265-273_ peptide. (**B**) Representative dot plots of bulk (cell dye-negative) and IGRP_265-273_ directed (cell dye-positive) CD8^+^ T cells incubated for 20 h with the indicated stimuli using K562/A*0201 cells for antigen presentation in the dye-based CD8^+^ T cell activation assay. Cells in the CD8 gate are shown. (**C**) t-SNE plots of single-cell gene expression data from IGRP_265-273_-directed CD8^+^ T cells after incubation for 20 h with the indicated antigen-presenting cells in the presence of solvent (black), mock peptide (blue), or cognate peptide (red). (**D**) Heatmaps of single-cell gene expression data for the previously identified 16 genes of antigen-responsive virus-directed CD8^+^ T cells derived from IGRP_265-273_-directed CD8^+^ T cells stimulated with the control or cognate peptide (see Fig. 3a). The reference genes *CD3E* and *CD8B* are also shown.

Thus, despite their strong binding to IGRP peptide-loaded multimers and distinct TCR repertoire, the antigen responsiveness of these antigen-specific cells is not analogous to that of virus-directed cells when tested in the same stimulatory setting. These findings suggest that certain autoantigen-specific CD8^+^ T cells show differing responses to their cognate peptide, are anergic, or may need divergent stimulatory conditions to mount a response similar to that of virus-specific CD8^+^ T cells.

### Discussion

The use of multimers to identify and subsequently characterize antigen-specific CD8^+^ T cells has advanced our knowledge of human TCRs and their phenotype. Here, we demonstrated that multimer-based identification of antigen-specific CD8^+^ T cells is adversely affected by recent activation, and we identified large gene sets that were congruently up- or downregulated in virus antigen-responsive CD8^+^ T cells across multiple viruses and donors and under different conditions of antigen presentation. We developed algorithms that accurately identify virus-responsive CD8^+^ T cells and can be applied broadly to single-cell RNA profiling data from antigen-stimulated cells.

Our approach to defining antigen-responsive profiles included proven models of well-established viral peptides presented by HLA-A*0201. We used two antigen presentation schemes and dominant peptides from two viruses. The antigen-specific cells that were used to establish the profiles were highly specific, as judged by their strong multimer binding and verified by TCR sequencing. Finally, we demonstrated a broad methodological applicability using three RNA profiling methods, and chose genes that identified virtually all of the multimer-selected antigen-responsive cells.

The limitations of this approach include a restricted set of antigenic peptides, which prevents us from generalizing our findings. In particular, the responsive profiles may be inappropriate for non-viral peptides, as we demonstrated for the islet β cell autoantigen IGRP. It is possible that the multimer-selected cells did not cover all phenotypes of antigen-specific CD8^+^ T cells, including cells with anergic or exhausted profiles. About 1% of the multimer-sorted virus-specific cells were not activated after cognate peptide presentation, and these cells could represent anergic or other T cell types. Finally, because our findings were based on *in vitro* stimulation, the profiles observed here may not represent those of CD8^+^ T cells activated *in vivo*.

Peptide stimulation of CD8^+^ T cells resulted in up- or downregulation of over 1000 genes. Of particular relevance was the downregulation of TCR on the cell surface, which was also associated with a marked reduction in multimer binding. This is consistent with previous findings (9, 11), suggesting that the frequency of antigen-specific CD8^+^ T cells may be underestimated in conditions of chronic or recent *in vivo* activation when using multimers for quantification. Notably, we observed a marked reduction in multimer staining, even after stimulation with the cognate peptide at a concentration of 1 ng/mL, which is 10,000 times lower than the concentration typically used for *in vitro* stimulation. A bias against selection of recent or chronically activated CD8^+^ T cells by multimers may grossly underestimate the degree of CD8^+^ T cell activation in disease, and could confound recent findings in diseases such as type 1 diabetes (24).

Our objective was to identify a panel of genes and a classifier that would enable us to identify *in vitro*-stimulated, antigen-responsive CD8^+^ T cells from unselected populations. This requires high specificity because the antigen-responsive cells comprise a minority of cells. High specificity would be provided by genes that are consistently differentially expressed with high quantification in responsive cells. A set of 10 genes, identified by different methods, was used to develop a classifier using machine learning methods. The approach was used to identify antigen-responsive CD8^+^ T cells from single-cell data, and provides a tool for the future refinement of algorithms suitable for other virus antigen-responsive and other antigen-responsive cells.

At the protein level, we observed increased expression of identified markers CD82 and CRTAM (CD355), similar to the commonly used activation markers CD137 and CD25, in antigen-responsive cells. Our study found many other genes that are known to be transcriptionally increased or that encode proteins that are increased in antigen-responsive CD8^+^ T cells. These include XCL1 and XCL2 (8, 15, 25, 26), EGR2 (27, 28), NR4A2 (8), and MIR155HG (29, 30), among other proteins (31–33). Some of the identified genes encode proteins that are often used to isolate antigen-specific cells. IFNγ, which is used for cytokine capture of responsive CD8^+^ T cells (34), was strongly upregulated in the majority of the activated cells. However, it was noticeably absent in a proportion of the antigen-responsive cells, indicating that such isolation methods may miss some responsive cells. We also identified several genes that were differentially expressed in the antigen-responsive CD8^+^ T cells and, to our knowledge, have not been previously described in this context. These include genes encoding the transcription factor ZBED2, which regulates transcription by RNA polymerase II, and was exclusively upregulated in many of the antigen-responsive cells; the guanine nucleotide binding plasma membrane associated protein GNG4; HSPA8; and the ribosomal protein RPLP0. These findings may help to understand the processes involved in the T cell-mediated antiviral response and provide new therapeutic targets.

A practical aspect of our study is that we accurately traced the virus-responsive cells identified in the transcriptomic profile back to their TCRs. This allowed us to perform broader antigen screening in combination with large-scale scRNAseq, where the responsive transcriptomic profile is used to map the antigen-specific TCR. An advantage of this approach is that the TCRs are representative of cells that can be robustly activated and permits selection of cells expressing desired effector molecules. This may be useful for vaccine development. Although we have not shown that similar profiles can identify tumor antigen-responsive CD8^+^ T cells, it is likely that the identification of TCRs using a similar approach may facilitate the development of chimeric antigen receptor T cell-based therapies that are patient-specific. Of potential interest, we found no signature that could identify cells responsive to a known epitope of a β cell autoantigen, despite the high frequency of strong multimer-positive memory CD8^+^ T cells in the donor. We previously reported that IGRP-directed clones generated from the same donor (and expressing the identical TCR) can kill peptide-loaded target cells *in vitro*, albeit with delayed kinetics (23). This suggests the possibility of an antigen-directed response of IGRP-directed cells to the target, at least in certain *in vitro* conditions. It is unclear whether autoantigen-specific CD8^+^ T cells are generally less responsive than virus-specific cells or if our finding is limited to this antigen or this patient. A previous study showed that autoantigen-specific CD8^+^ T cells obtained from peripheral blood were less responsive than pancreatic cells (24). Further studies are needed in patients with active autoimmune disease.

In conclusion, we provide findings that will allow the identification of antigen-responsive CD8^+^ T cells and their TCRs in an unbiased manner. The findings can be applied to antigens where multimer reagents are unavailable, as an alternative to multimer-based cell selection or in combination with multimer selection. This should facilitate the development of CD8^+^ T cell activation-based therapies.

## Materials & Methods

### Subjects

Samples were obtained from six healthy adult blood donors or from an adult patient with type 1 diabetes; all had the HLA-A*0201 allele. All methods were performed in accordance with relevant guidelines and regulations. Samples were collected after obtaining informed consent and ethical committee approval (EK 240062016, TU Dresden and 5049/11, TU München).

### Peptides

The Flu MP_58-66_ (GILGFVFTL), hCMV pp65_495-503_ (NLVPMVATV) and IGRP_265-273_ (VLFGLGFAI) peptides were purchased from Mimotopes or Panatecs at >95% purity, as confirmed by high-performance liquid chromatography and mass spectrometry. For the T cell assays, the peptides were dissolved in dimethyl sulfoxide (DMSO) to a concentration of 10 mg/mL and subsequently diluted in assay medium to final peptide concentrations of ≤10 µg/mL. The DMSO concentration in each assay was ≤0.1%.

### Cell staining and flow cytometry

Cells were stained using the following surface marker-directed monoclonal antibody– fluorochrome combinations: CD3-APC (clone HIT3a), CD3-AF488 (clone UCHT1), CD4 (clone SK3), CD8-APC-Cy7 (clone SK1), CD14-APC (clone M5E2), CD19-APC (clone HIB19), CD56-APC (clone B159), CD25-PE-Cy7 (clone M-A251), CD82-AF647 (clone 423524), and CD95-CF594 (clone DX-2) from BD Pharmingen; CD137-PerCP-eFluor710 (clone 4B4-1) and CD355-APC (clone Cr24.1) from Thermo Fisher Scientific; and CD16-APC (clone 3G8) from BioLegend. 7-Aminoactinomycin D (7AAD, BD Pharmingen) or Sytox Blue (Thermo Fisher Scientific) was used to exclude dead cells. Phycoerythrin-labeled HLA-A*0201 multimers loaded with Flu MP_58-66_ (GILGFVFTL), hCMV pp65_495-503_ (NLVPMVATV), or IGRP_265-273_ (VLFGLGFAI) were purchased from Immudex. Cells were acquired on a flow sorter (BD FACSAria Fusion, BD Biosciences) with FACSDiva 8 software (BD Biosciences) and analyzed using FlowJo 10 software (FlowJo LLC). Unless otherwise stated, cells were stained in phosphate-buffered saline (PBS) supplemented with 1% pooled human AB serum (PBS/1%AB) for 30 min on ice followed by at least two washing steps using the same buffer, and were then stained with a marker for dead cells before analysis. For CFSE labeling, PBMCs (4 × 10^7^/mL) were resuspended in PBS at room temperature, mixed 1:1 with CFSE staining solution (1 µM CFSE in PBS), and the cell suspension was immediately vortexed for 10 s to achieve a homogenous CFSE distribution. Cells were incubated at 37 °C for 10 min in the dark and washed with X-VIVO15 media to remove excess CFSE. Multimer staining was carried out in PBS supplemented with 5% pooled AB serum for 10 min at room temperature followed by 25 min incubation on ice in the presence of the respective antibodies. Cells were subsequently washed with PBS/1%AB and incubated with 7AAD or Sytox Blue for 10 min immediately before flow cytometry. The gating strategies used for the analysis and sorting of multimer- or CFSE-stained cells, and for the isolation of antigen-responsive cells and CD95^+^ CD8^+^ memory T cells are illustrated in Fig. S8.

### CD8^+^ T cell activation assay

The activation assays were conducted using 200–500 CFSE-labeled antigen-specific CD8^+^ T cells together with either 5 × 10^5^ unlabeled PBMCs (PBMC-based assays) or a combination of 2.5 × 10^5^ HLA-A*0201-expressing K562 cells (K562/A*0201 (35); kindly provided by Prof. Thomas Wölfel; Johannes Gutenberg Universität, Mainz, Germany) and 5 × 10^4^ flow-sorted bulk CD8^+^ T cells (cell line-based assays). To obtain CFSE-labeled multimer-specific CD8^+^ T cells, the PBMCs were labeled with CFSE and subsequently incubated with the respective peptide-loaded HLA-A*0201 multimers to identify and sort highly pure antigen-specific CD8^+^ T cells into the assay tubes. Unless stated otherwise, the cells were incubated with the cognate peptide (10 µg/mL), mock peptide (10 µg/mL), or solvent (DMSO) for 18–24 h.

Subsequently, viable CFSE^+^ CD8^+^ T cells were single-cell-sorted for downstream gene expression analysis and TCR sequencing. For some experiments, antigen-responsive cells were sorted based on their expression of the activation marker CD137. In these cases, the assays were performed without prior CFSE labeling or multimer staining.

For droplet encapsulation sequencing experiments conducted on the 10x Genomics platform, PBMCs were incubated with Flu MP_58-66_ or hCMVpp65_495-503_ peptide for 20 h and 6500 viable CD95^+^CD8^+^ memory CD8^+^ T cells were sorted from each of these conditions and processed for gene expression and TCR analysis.

### Targeted single-cell gene expression analysis

Targeted gene expression analysis was done by single-cell qPCR as previously described (23) with some modifications. cDNA was synthesized using Quanta qScript TM cDNA Supermix directly on cells. Total cDNA was pre-amplified for 20 cycles (1 × 95 °C for 8 min, 95 °C for 45 s, 49 °C with 0.3 °C increment/cycle for 1 min, and 72 °C for 1.5 min) and 1 × 72 °C for 7 min with TATAA GrandMaster Mix (TATAA Biocenter) in a final volume of 35 μL in the presence of the 75 primer pairs listed in Table S9. The final concentration of each primer was 25 nM. Pre-amplified cDNA (10 μL) was then treated with 1.2 U exonuclease I and expression was quantified by real-time PCR on a BioMarkTM HD System (Fluidigm Corporation) using the 96.96 Dynamic Array IFC, GE 96 × 96 Fast PCR+ Melt protocol, and SsoFast EvaGreen Supermix with Low ROX (Bio-Rad). The primer concentration was 5 μM in each assay.

### Single-cell RNA sequencing

The scRNAseq workflow was based on the previously described Smart-seq2 protocol (36) with the following modifications. Single cells were flow-sorted into 96-well plates containing 2 µL of nuclease-free water with 0.2% Triton X-100 and 4 U murine RNase inhibitor (NEB), centrifuged, and frozen at −80 °C. After thawing, 2 µL of the primer mix (5 mM dNTP [Invitrogen], 0.5 µM oligo-dT primer, 4 U murine RNase inhibitor) was added to each well. The reverse transcription reaction was performed as described (36), but with final concentrations of RNase inhibitor and Superscript II of 9 U and 90 U, respectively, at 42 °C for 90 min, followed by an inactivation step at 70 °C for 15 min. The number of pre-amplification PCR cycles was increased to 22 to ensure there was sufficient cDNA for downstream analysis. The amplified cDNA was purified using 1× Sera-Mag SpeedBeads (GE Healthcare), and DNA was eluted in 12 µL nuclease-free water. The concentration of samples was measured using a plate reader (Infinite 200 PRO, Tecan) in 384 well black, flat-bottom, low-volume plates (Corning) using an AccuBlue Broad Range kit (Biotium). Then, 0.7 ng of pre-amplified cDNA was used for library preparation (Nextera DNA library preparation kit, Illumina) in a 5-µL reaction volume. Illumina indices were added during the PCR reaction (72 °C for 3 min, 98 °C for 30 s, 12 cycles of [98 °C for 10 s, 63 °C for 20 s, and 72 °C for 1 min], and 72 °C 5 min) with 1× KAPA Hifi HotStart Ready Mix and 0.7 µM of dual indexing primers. After PCR, the libraries were quantified with AccuBlue Broad Range kit, pooled in equimolar amounts, and purified twice with 1× Sera-Mag SpeedBeads. The libraries were sequenced on the NextSeq 500 Illumina platform to obtain 75-bp single-end reads aiming at an average sequencing depth of 0.5 million reads per cell.

### Droplet encapsulation sequencing

The scRNAseq was performed on the 10x Genomics platform using Chromium Single Cell Immune profiling 5’ v2 Reagent Kits according to the manufacturer’s instructions. Gene expression libraries were sequenced on the NextSeq 500 Illumina platform using a high-output flow cell to obtain paired-end reads at the following read lengths to generate ∼160–180 million fragments: read 1, 26 cycles, i7 index, 8 cycles; read 2, 57 cycles. Similarly, TCR-enriched libraries were sequenced on the MiSeq Illumina platform to obtain 150-bp paired-end reads at a depth of 15 million total read pairs.

### Data processing and analysis

#### Targeted single-cell gene expression analysis

Raw data were analyzed using Fluidigm Real-Time PCR analysis software. Preprocessing and data analysis were conducted using KNIME 2.11.2 and RStudio Version 0.99.486. Preprocessing with a linear model to correct for confounding factors was conducted as previously described (37). To model the bimodal gene expression of single cells from T cell clones, the Hurdle model, a semi-continuous modelling framework, was applied to the preprocessed data (38). This allowed us to assess the differential expression profiles with respect to the frequency of expression and the positive expression mean via a likelihood ratio test. Genes were ranked by a score derived from the number of comparisons with statistically significant differences in gene expression and the number of donors with statistically significant comparisons (0.5 points per significant comparison multiplied by the number of donors showing significant differences in expression of each gene). Genes with the same score were ranked based on the delta of their median expression differences between the cognate and control stimuli.

#### Single-cell RNA-sequencing

The scRNAseq adapter trimmed reads were mapped against the human hg38 reference genome using STAR (v2.5.4, default parameters) and Ensembl genes (v 91) as the gene model references for alignment to produce one BAM file per cell. The count per gene matrix was obtained using featureCounts (v1.6.2, *-Q 30, -p*). Differentially expressed genes were identified by performing pairwise comparison using two independent methods: SCDE (v2.8.0) (39) and DESeq2 (v1.20.0, using a zero-inflated negative binomial distribution) (40). Low-quality cells expressing only a few genes were filtered out from the counts matrix using the *clean.counts* function in SCDE (*min.lib.size* = 1000, *min.reads* = 1, *min.detected* = 1). Only genes that were identified as differentially expressed by both tools were considered to be expressed differentially. Data from each experiment were visualized using the t-SNE algorithm implemented in the Rtsne R package. In all comparisons, the cells stimulated with cognate peptide were labeled as the test group, while cells incubated with solvent or mock peptide showed similar profiles, and thus were combined as a control group. Because the SCDE method does not output a *p*-value for each differentially expressed gene, a two-sided *p*-value for each gene was calculated from the Benjamini–Hochberg multiple testing corrected Z score (cZ) using the normal distribution as the null hypothesis. Genes were ranked by the product of their log_2_-fold change and the reciprocal of the adjusted *p*-value (values obtained from SCDE). For experiments using several donors, the genes were ranked individually for each donor and a composite list was generated by obtaining a mean rank for each gene across the multiple donors and then ranking the genes based on their mean rank. To identify genes that could distinguish cognate peptide-stimulated cells and control-stimulated cells in individual experiments, we calculated the separation index (SI) for each gene (*g*) using the following equations.

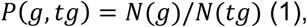

where *tg* is the test group, and *N* is the number of cells. To further clarify, *N(g)* is the number of cells with a non-zero count for the particular gene, and *N(tg)* is a total number of cells in the test group. The same applies to the other groups analyzed.

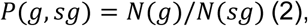

where *sg* is the solvent group.

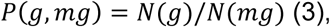

where *mg* is the mock group.

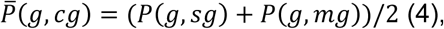

where *cg* is the control group.

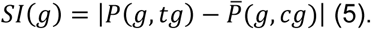

The genes were ranked according to their SI in each experiment in descending order, and then ordered by the mean rank of all PBMC-based assays. For the top 50 genes, we determined the median expression difference between cognate peptide- and control-stimulated cells to identify genes that were potentially usable as marker genes.

The Enrichr package was used to determine whether the differentially expressed genes were preferentially enriched in a particular Kyoto Encyclopedia of Genes and Genomes (KEGG) pathway. To identify potential pathways, we performed pathway enrichment analysis using the following workflow. First, all pathways that showed enrichment (adjusted *p*-value < 0.05) in at least one of the four donors (three CMV and one Flu) were listed together to generate a unique list of these pathways. Then, for each of these pathways, the coverage index was calculated for each donor as the number of differentially expressed genes belonging to a pathway divided by the total number of genes in this pathway. Finally, the pathways were ranked by the median coverage index for each pathway across the four donors.

#### Droplet encapsulation sequencing

The raw gene sequencing data were processed using the *count* command in Cell Ranger software (v2.1.0, *--expect-cells=3000*) (10x Genomics). Gene annotation was filtered using the *mkgtf* command to include only protein-coding, lincRNA, and antisense gene features (*--attribute=gene_biotype:protein_coding, --attribute=gene_biotype:lincRNA, --attribute=gene_biotype:antisense*). The Seurat package (41) was used to analyze the gene expression data obtained in droplet encapsulation sequencing experiments. Briefly, all genes that were expressed in at least three cells and all cells that expressed ≥200 such genes were retained. The cells were then clustered based on the first 14 principal components (resolution parameter set to 0.6) and visualized using the resulting t-SNE (yielding 8 clusters) and combined with the TCR information retrieved from TCR-enriched libraries of the same experiment performed in parallel. CMVpp65_495-503_-directed cells were identified within the CD95^+^CD8^+^ memory T cell single-cell data via their TCRα and β chain sequences (using TCR information of CMVpp65_495-503_-directed cells retrieved in the preceding multimer-based scRNAseq experiments of the donor). To identify marker genes that separated the main cluster containing CMV-directed cells after stimulation with the cognate peptide from the remaining clusters, we used the *FindMarkers* function (test parameter set to *roc*). The top 10 marker genes for this main cluster were used to build a support-vector machine (SVM)-based classifier to discriminate between responsive cells and non-responsive cells. For this purpose, we used the SVM implementation available with the package libSVM (v1.04). The log-normalized counts for the 10 marker genes were exported to a file, and responsive cells were labeled as true positives, whereas all other cells were labeled as true negatives. This dataset was used to build the model, and the data obtained from the experiment with Flu MP_58-66_-stimulated bulk-sorted CD95^+^CD8^+^ memory T cells was used as the test set. The model was also used to identify responsive cells obtained from other scRNAseq-based experiments. For this, the experiment read counts were normalized using the DESeq2 *counts* function (with normalized = TRUE parameter), and then used as test sets.

### TCR identification

TCR sequences were reconstructed from the transcriptome of each cell using the TraCeR algorithm (v0.6.0, default parameters) (42). Reads originating from the mRNA of the TCRs were extracted from the sequence data for each cell and were assembled into contigs to generate full-length TCRs. Only productive TCR chains, both α and β, were retained for each cell. To resolve instances where one cell had more than one productive α or β chain, the chain with greater expression (obtained by parsing the transcripts per million count) was retained as the productive chain for the respective cell. Sequences of productive chains were then uploaded to IMGT/HighV-QUEST (43) to identify the V, D, and J genes as well as the CDR3 region associated with each chain. The TCR sequences of the droplet encapsulation experiments were extracted using Cell Ranger software.

### Additional statistical analyses

Wilcoxon’s matched-pairs signed-rank test was used to compare median fluorescence intensities for the analyzed surface markers among antigen-specific and antigen-responsive CD8^+^ T cells.

Software: FACS Diva 8, FlowJo 10, Graph Pad Prism 7, Tracer (v0.6.0), STAR (v2.5.4) libSVM (v1.04), featureCounts (v1.6.2), and R (v3.5.1) R-packages: DESeq2 (v1.20.0), SCDE (v2.8.0), EnrichR (v1.0), Seurat (v2.3.4), Rtsne (v0.13), pheatmap (v1.0.10), and ggplot2 (v3.1.0)

## Acknowledgements

We thank the CMCB flow cytometry core facility, particularly A. Gompf and K. Bernhardt for technical support, and L. Ventola for administrative support.

## Author contributions

Y.F.F., V.S., and E.B. designed the studies. Y.F.F., G.K., R.M., S.R., A.L., and D.L. performed the experiments. Y.F.F., A.E, A.D., and E.B. supervised the work. V.S., Y.F.F., A.P. and A.E. analyzed the data. Y.F.F., V.S, G.K., and E.B. wrote and edited the manuscript.

## Supplementary Materials

**Fig. S1:**
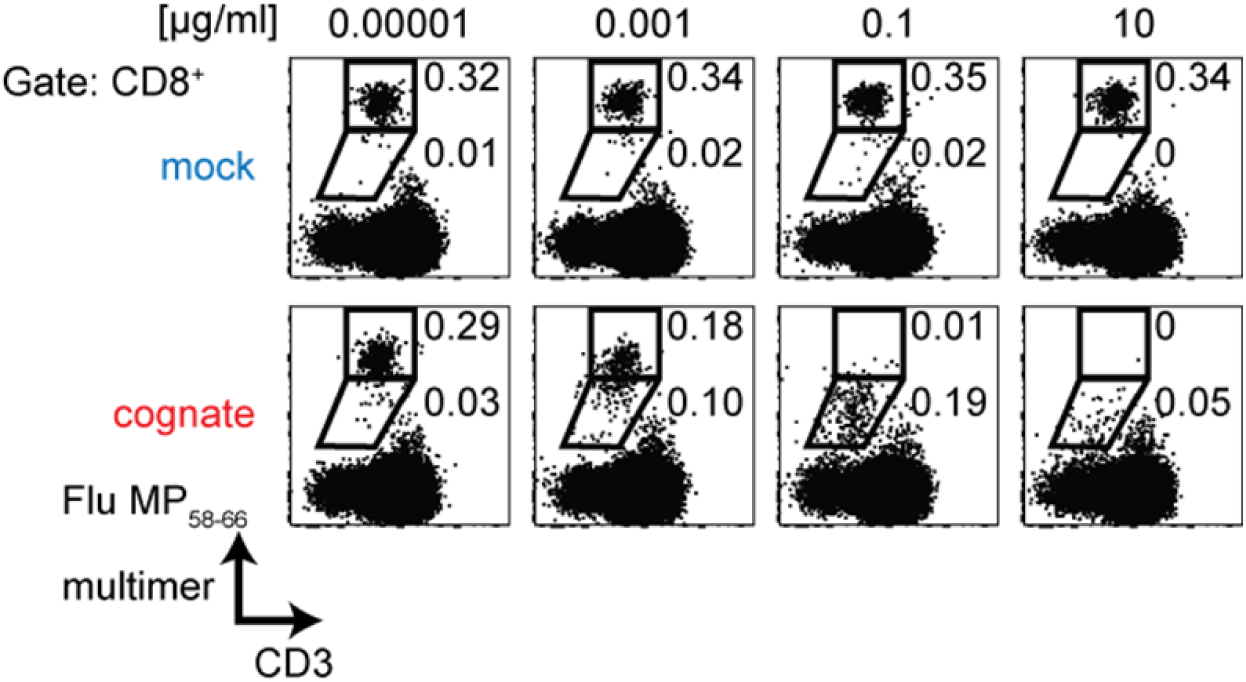
Impaired MHC class I multimer detection of CD8^+^ T cells responding to their cognate antigen. Representative flow cytometry dot plots of PBMCs stained with HLA-A2 multimers loaded with Flu MP_58-66_ peptide after incubating PBMCs for 20 h in the presence of mock (IGRP_265-273_) or cognate (Flu MP_58-66_) peptide at the indicated concentrations. Plots show 1 × 10^5^ cells in the CD8 gate and the frequencies of cells showing strong or intermediate positive staining with the multimer.

**Fig. S2:**
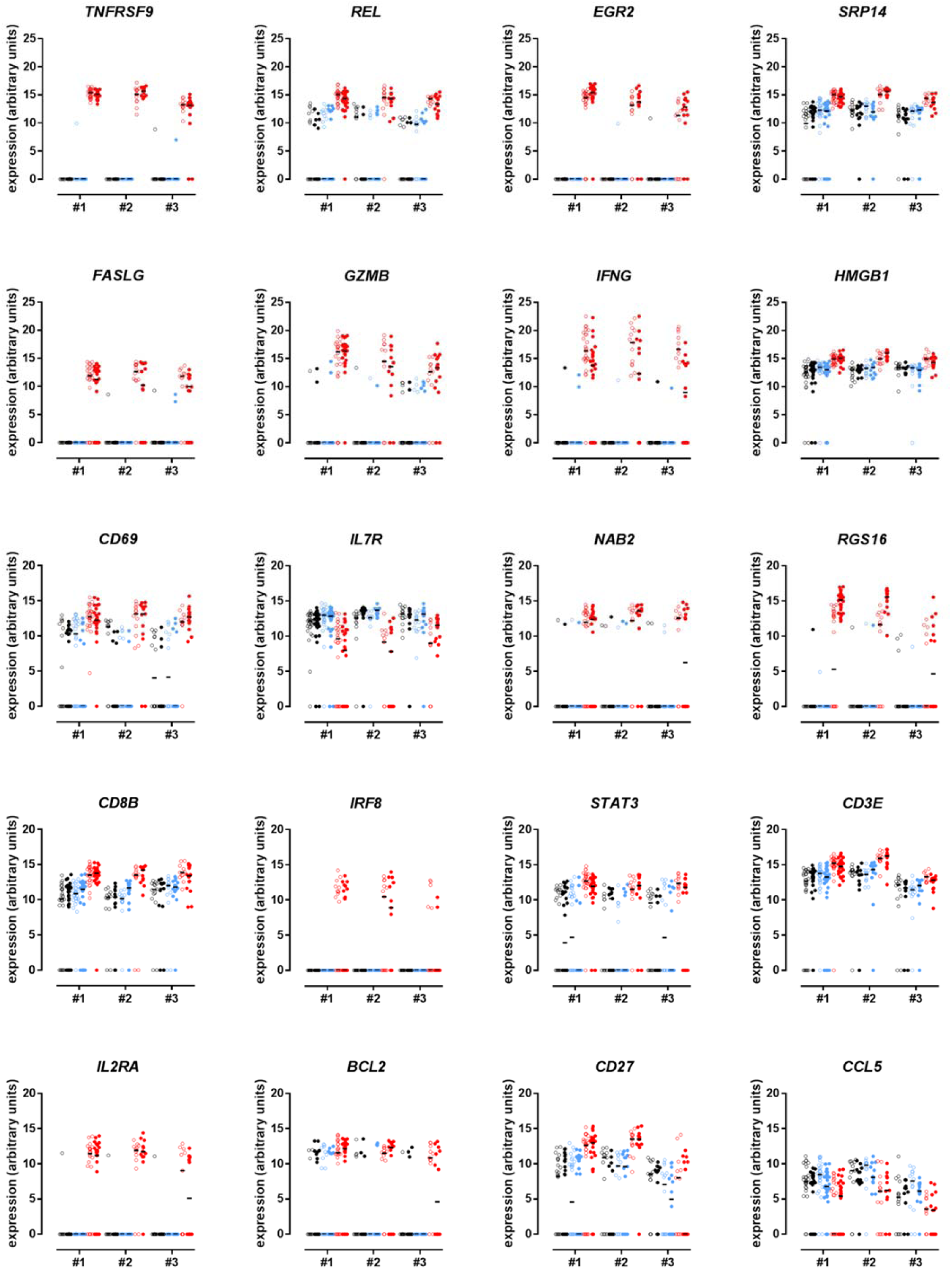
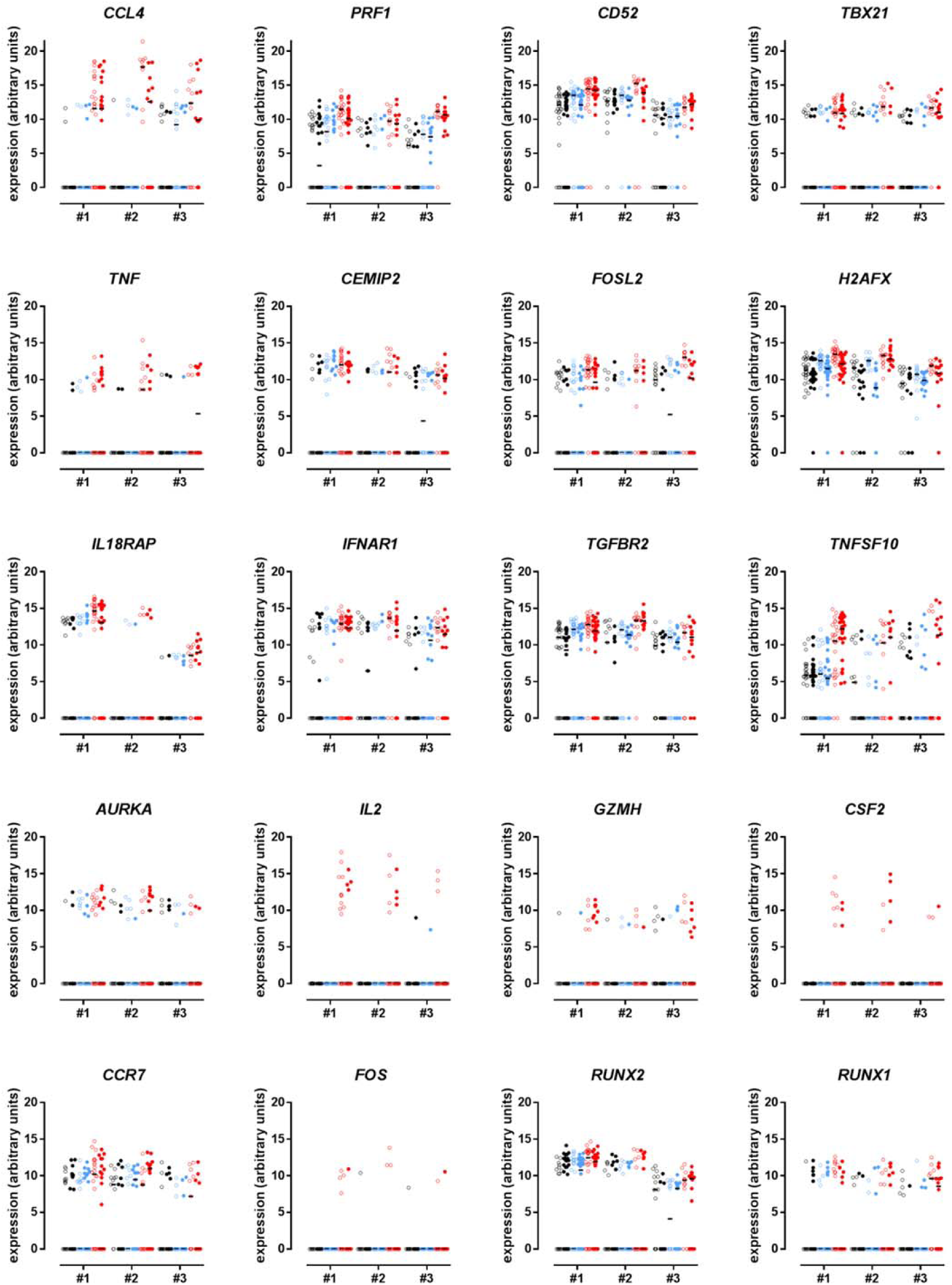
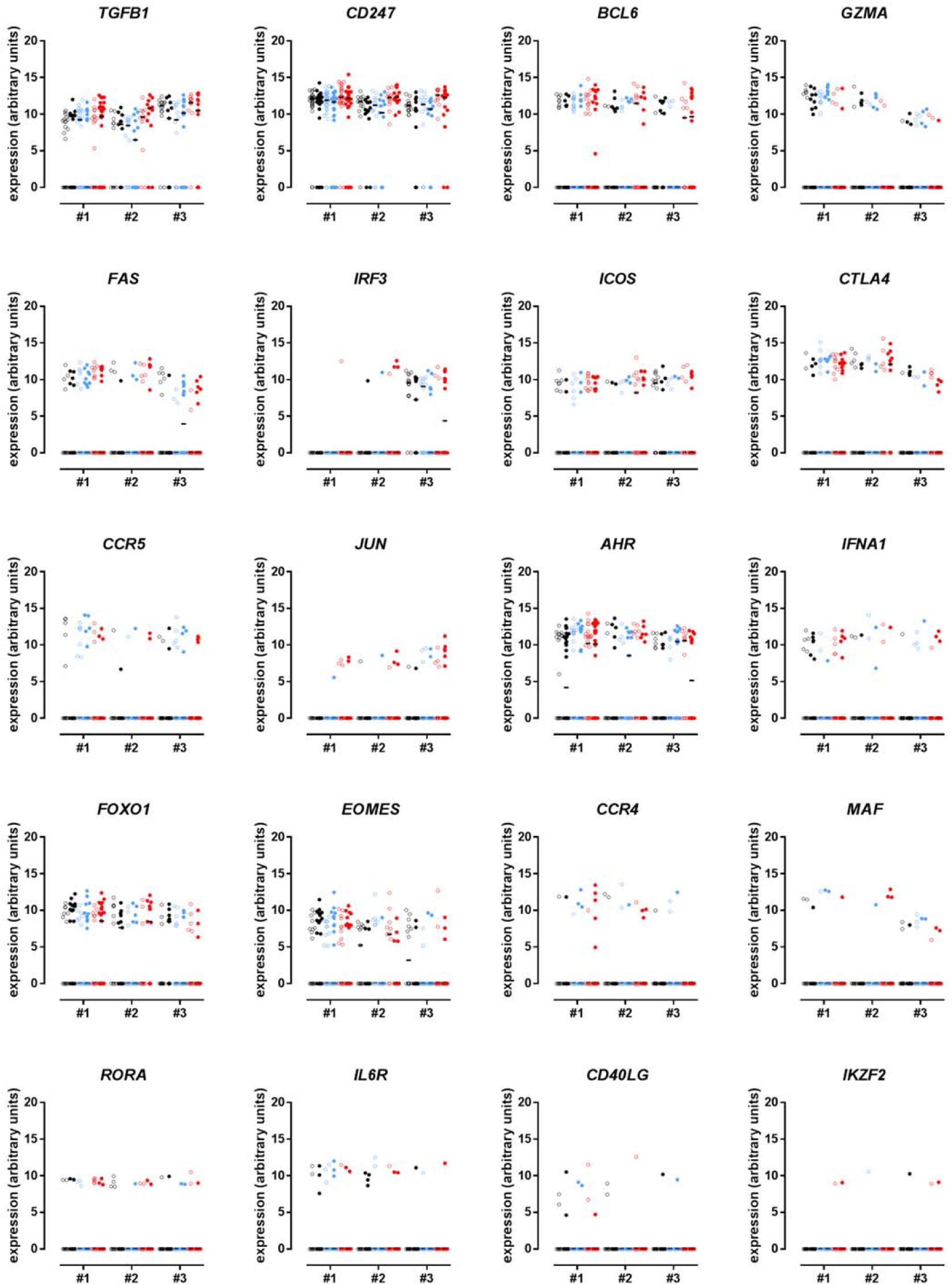
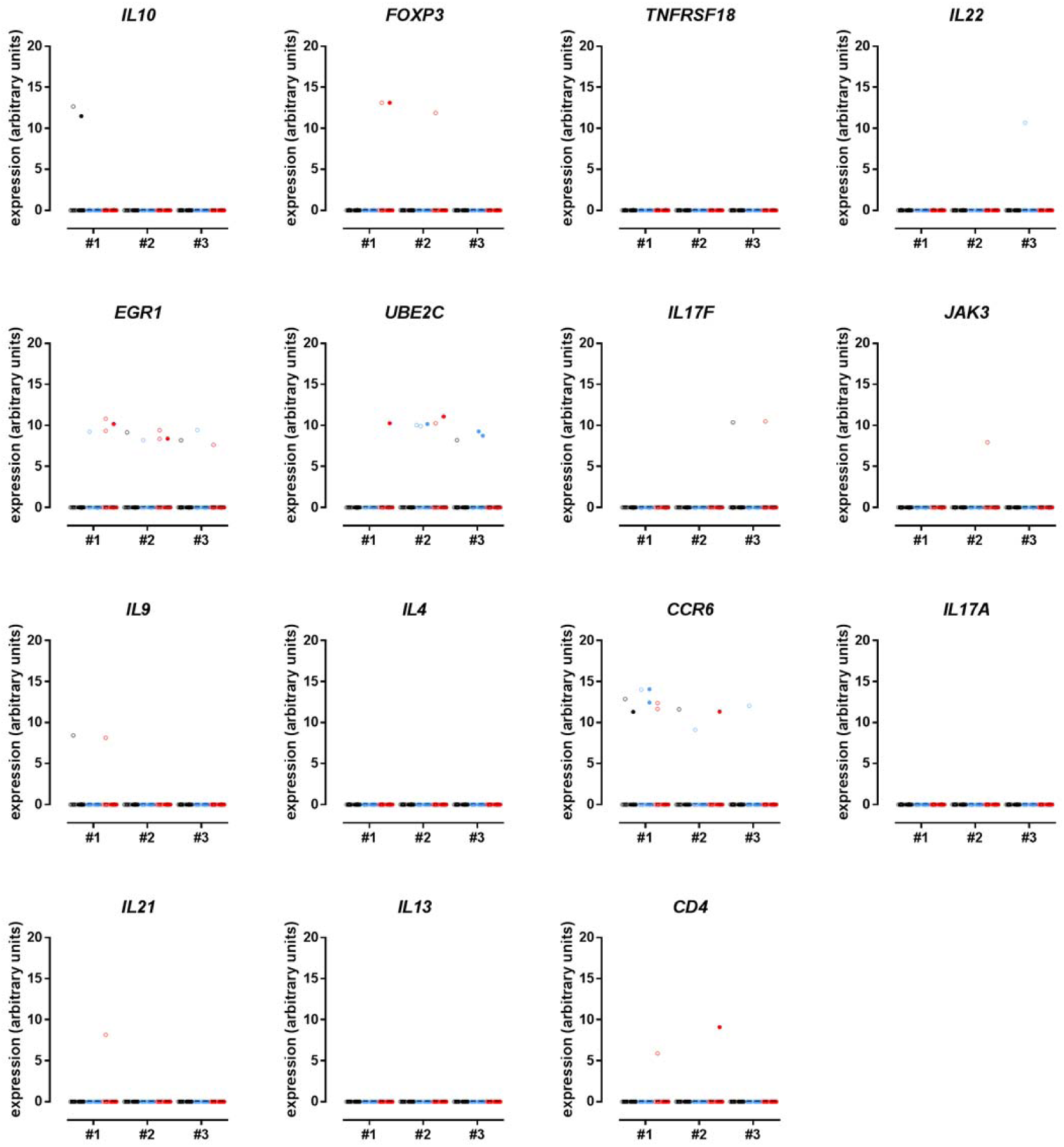
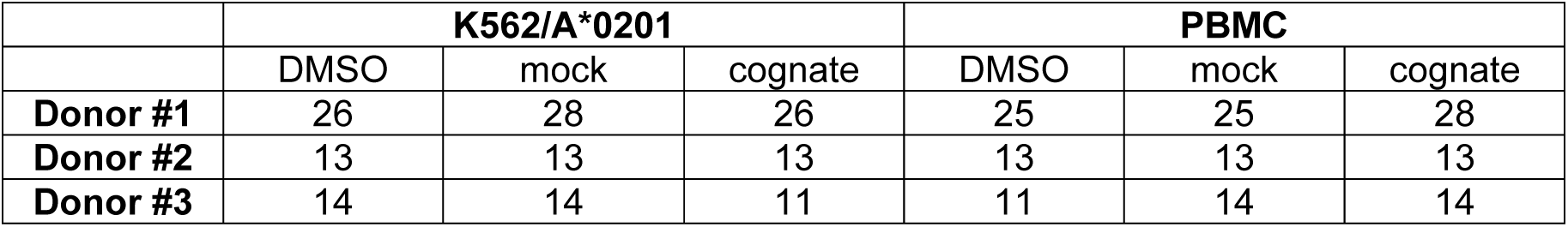
Gene expression analysis of Flu MP58-66-directed CD8+ T cells after incubation with peptide-loaded K562/A*0201 cells or autologous PBMCs. Scatter plots are shown for 75 genes determined using the qPCR-based targeted gene expression approach. Plots show the expression in individual Flu MP58-66-directed single-cell-sorted CD8+ T cells from donors #1–3 incubated overnight with DMSO (black) or mock peptide (blue) as control stimuli or with cognate peptide (red) in the presence of K562/*0201 cells (open symbols) or autologous PBMCs (filled symbols) for antigen presentation in the dye-based CD8+ T cell activation assay. The qPCR primers are listed in Table S9. The following numbers of cells were analyzed:

**Fig. S3:**
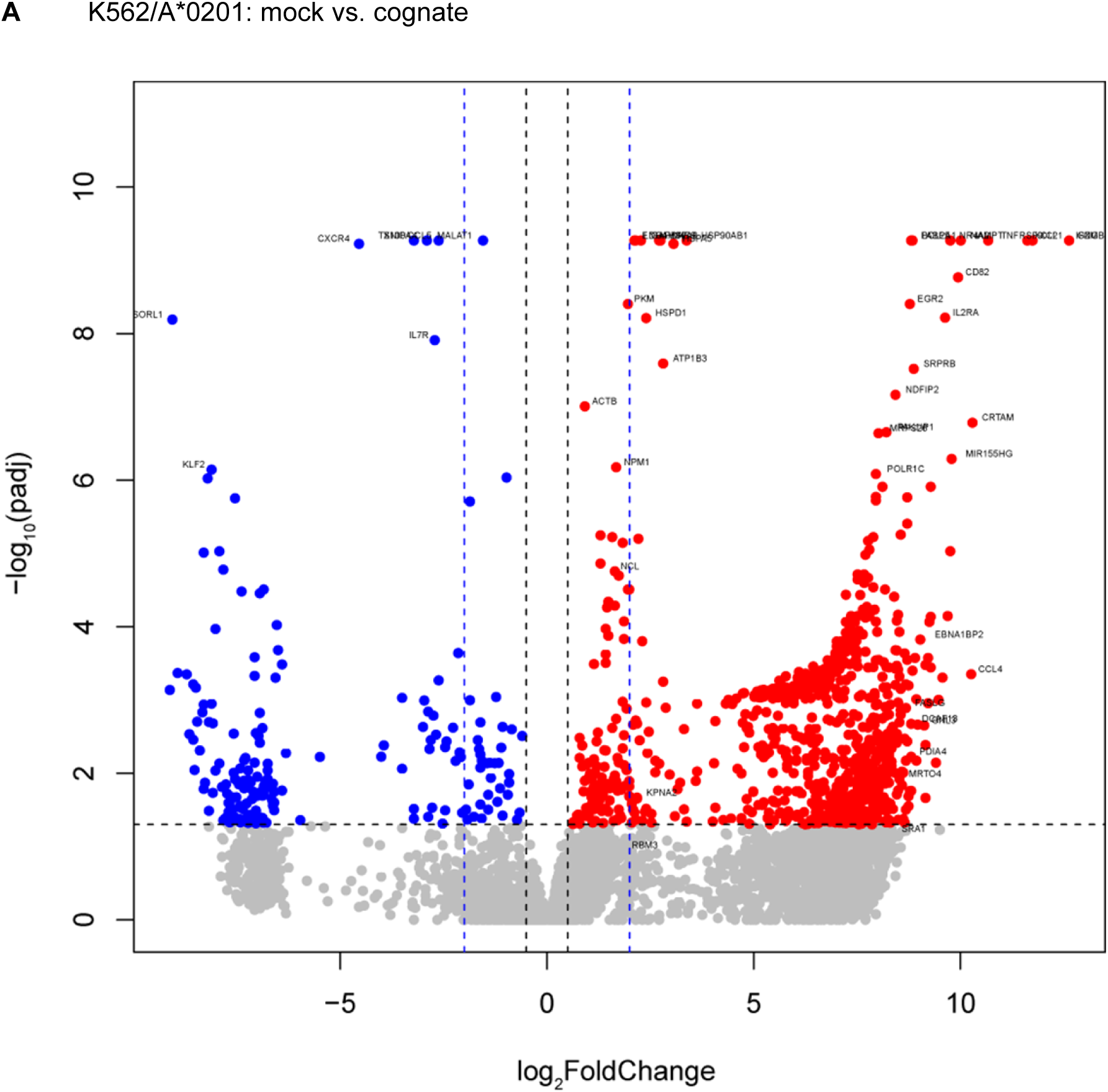
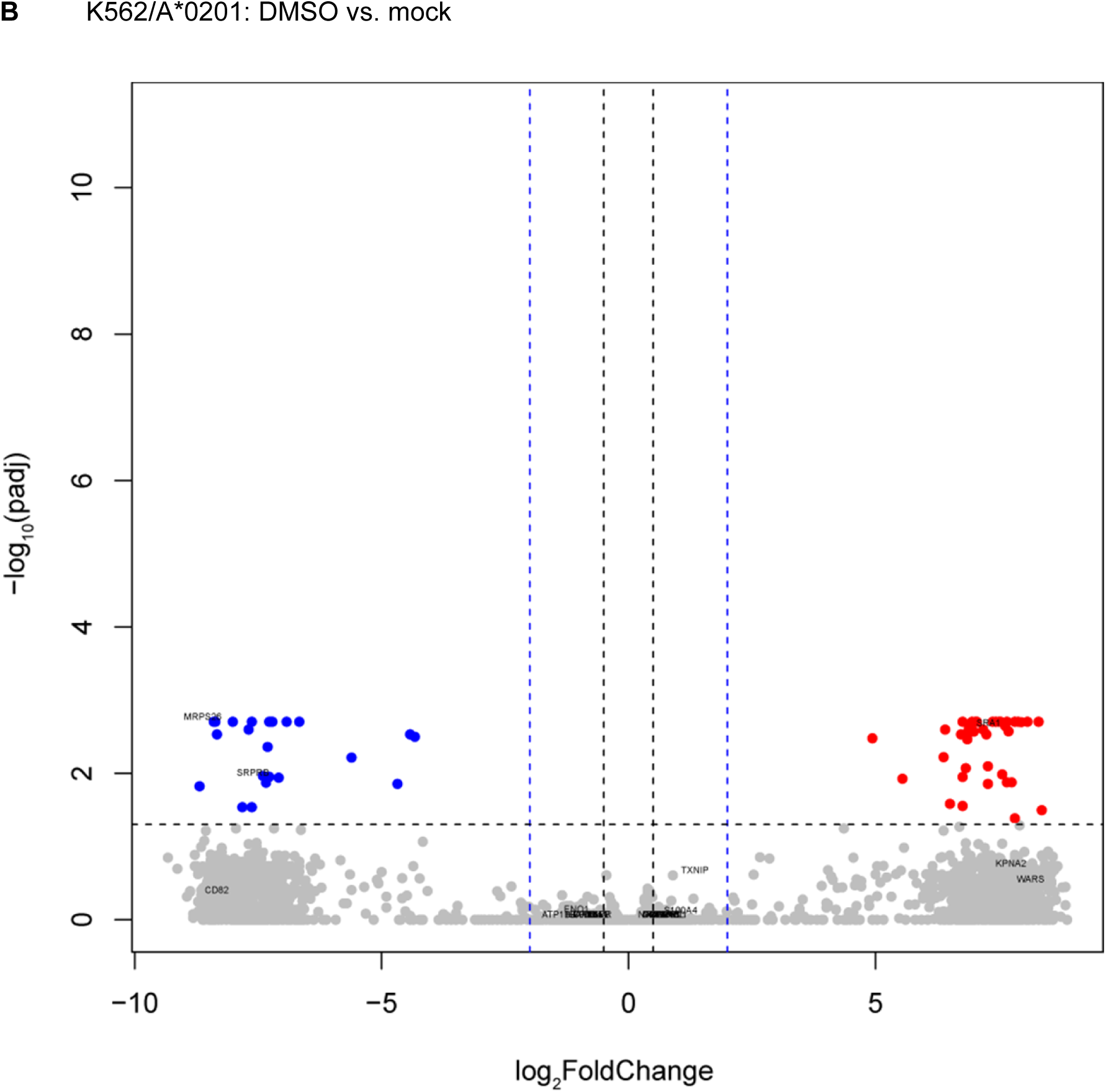
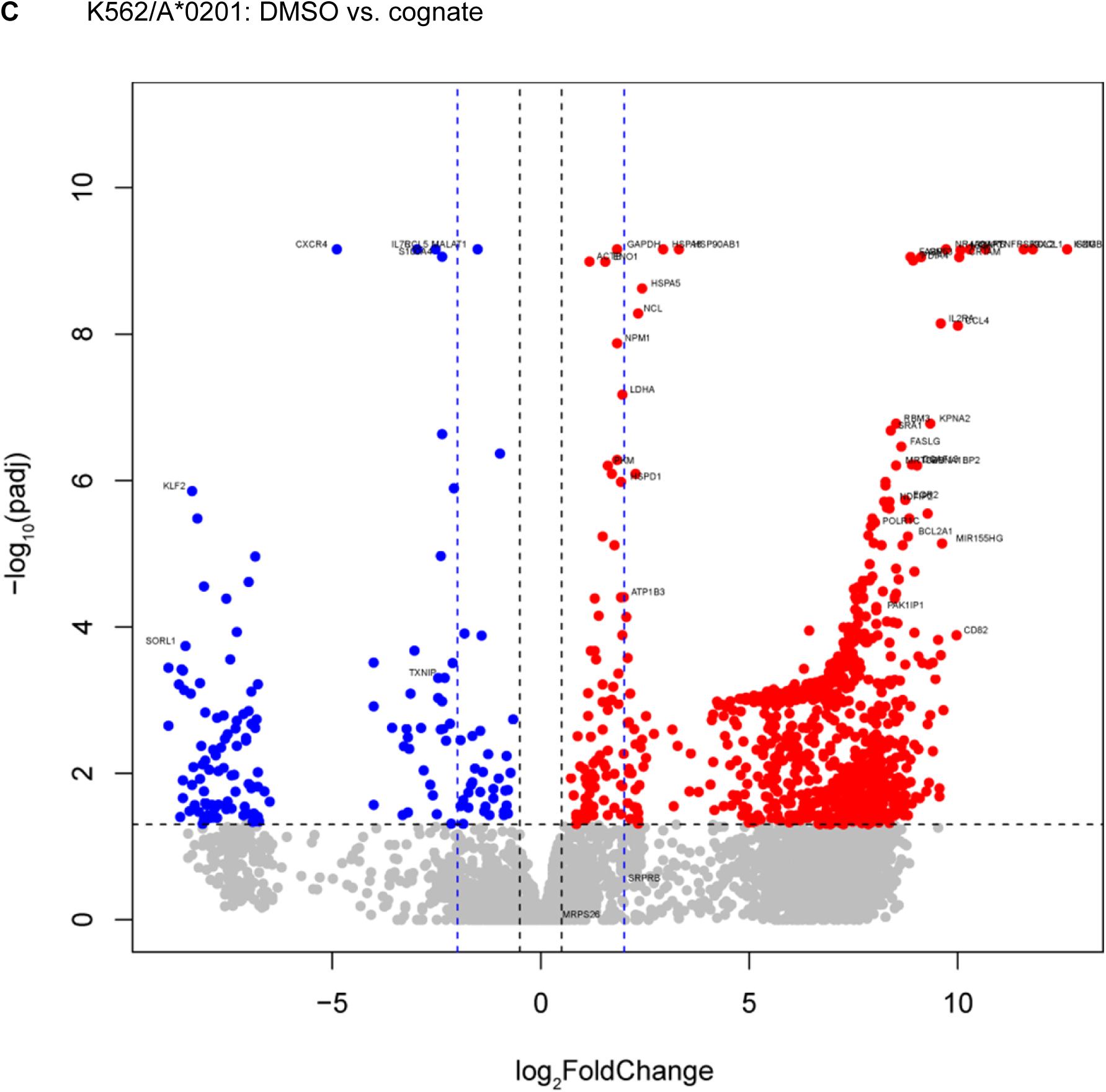
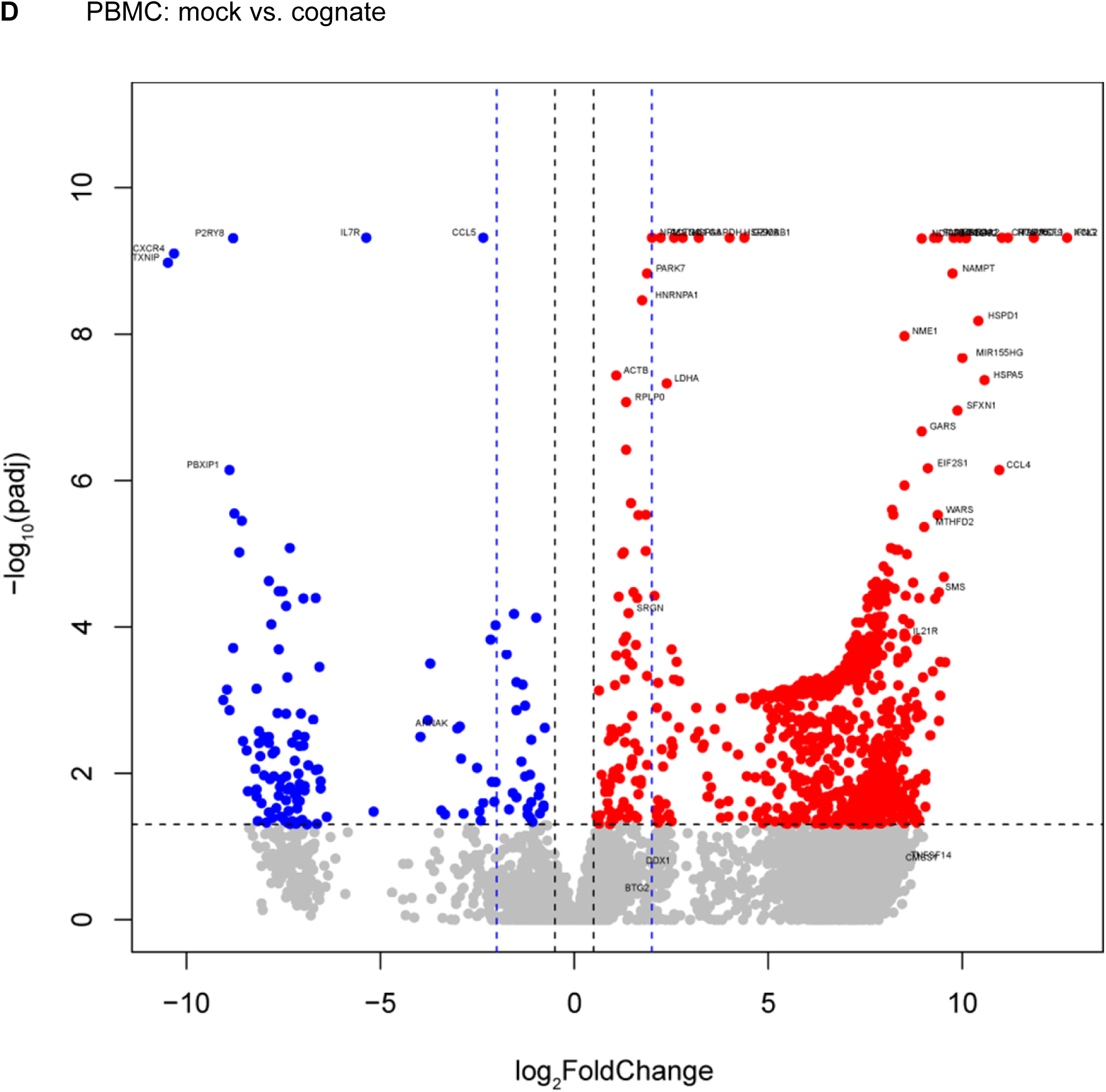
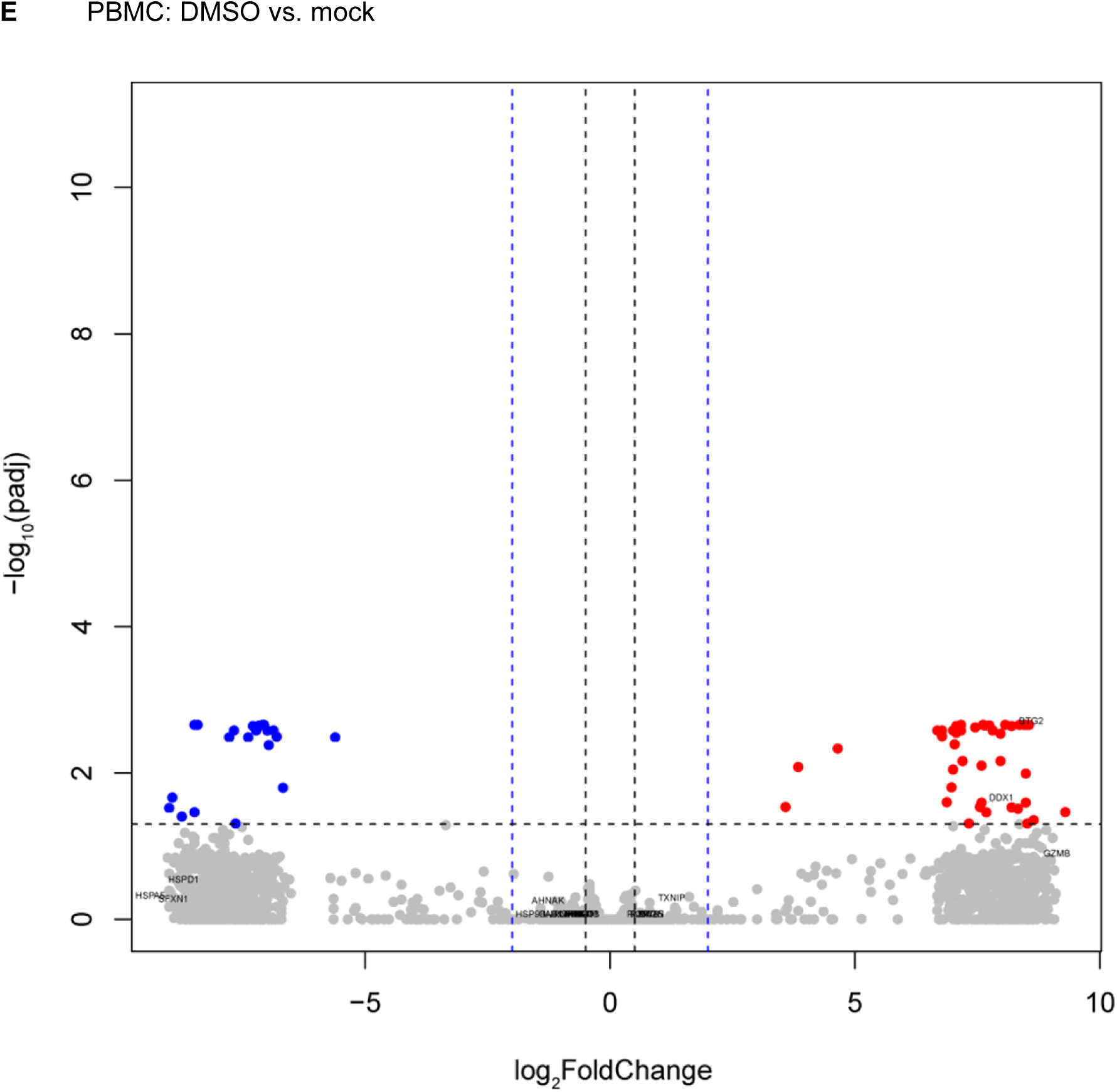
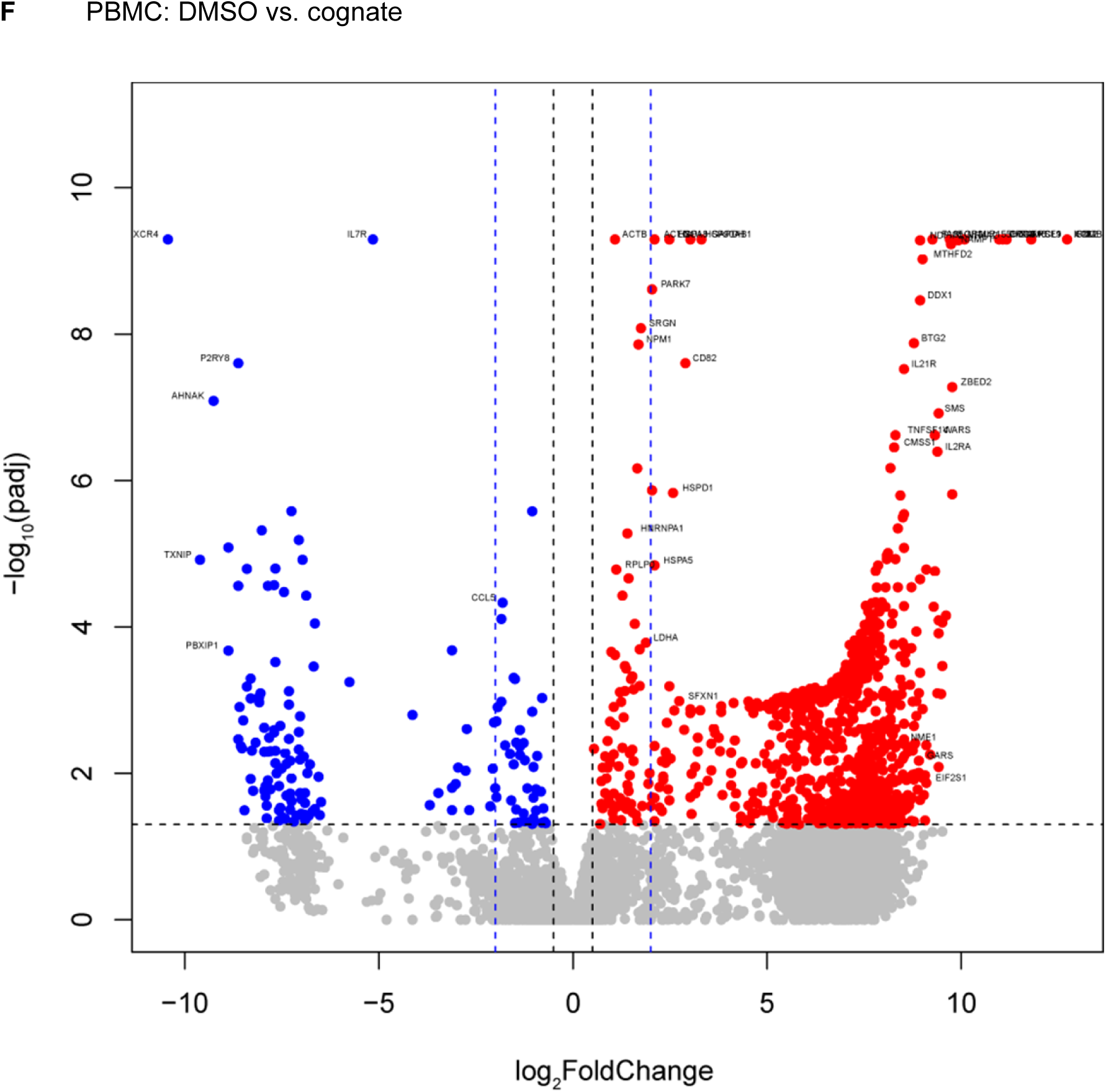
Comparison of gene expression between cognate peptide-stimulated and control-stimulated Flu MP_58-66_-directed CD8^+^ T cells. (**A–F**) Volcano plots of scRNAseq data for Flu MP_58-66_-directed cells from donor #1. Cells were single-cell-sorted from the dye-based CD8^+^ T cell activation assay following overnight incubation with peptide solvent (DMSO), mock peptide (IGRP_265-273_), or cognate peptide (Flu MP_58-66_) in the presence of K562/A*0201 cells or autologous PBMCs for antigen presentation. The thresholds (fold change = 0.5 [black] and 4 [blue]; adjusted *p* = 0.05 [black]) are shown as dashed lines. Genes upregulated in cells responding to their cognate peptide are marked in red and downregulated genes in blue. The gene symbols for the combined top 50 ranked differentially expressed genes in both comparisons (DMSO vs. cognate and mock vs. cognate) using the same cells for antigen presentation are shown. Volcano plots comparing cells incubated with peptide solvent and mock peptide (DMSO vs. mock) are also included as control conditions.

**Fig. S4:**
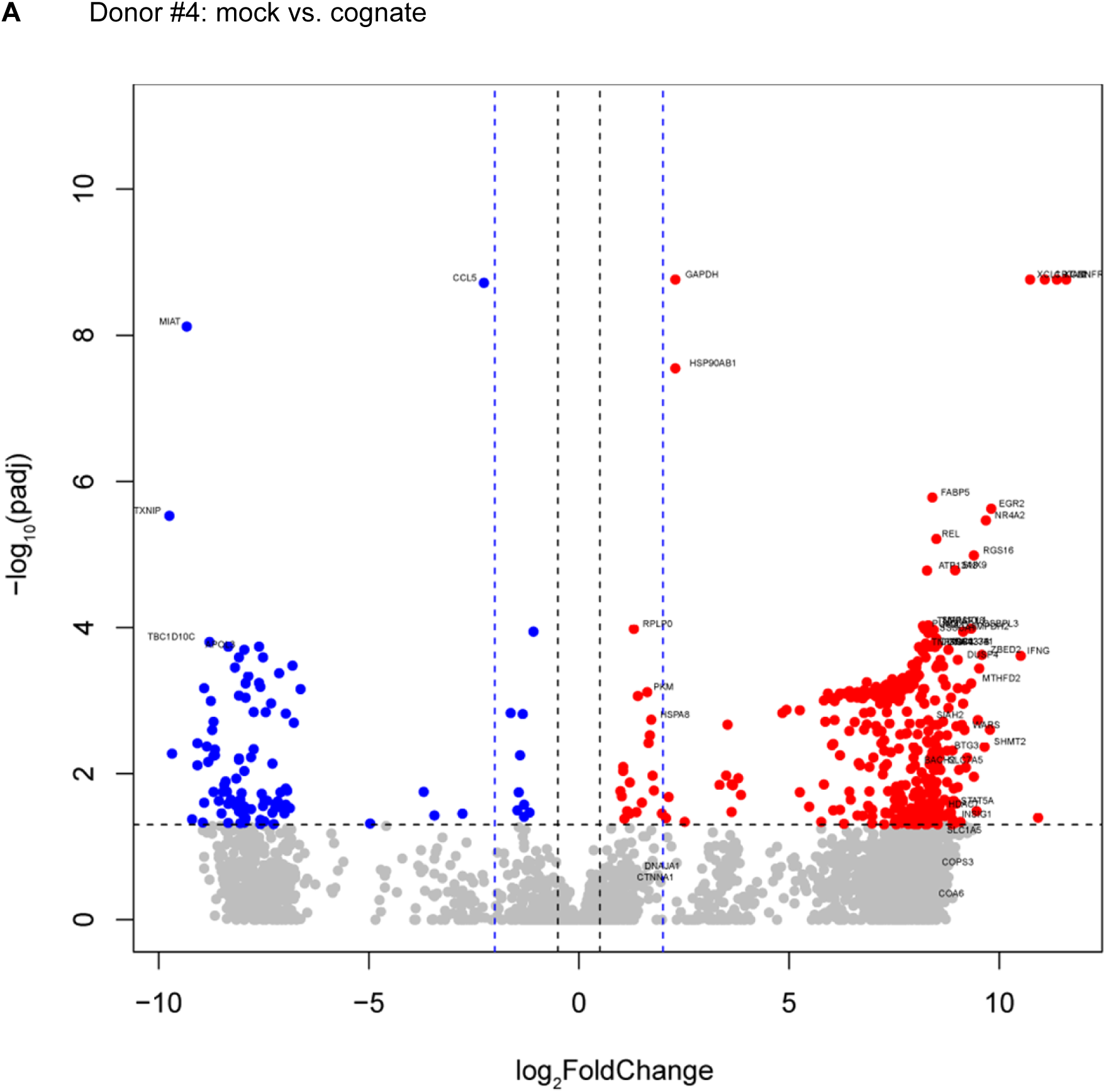
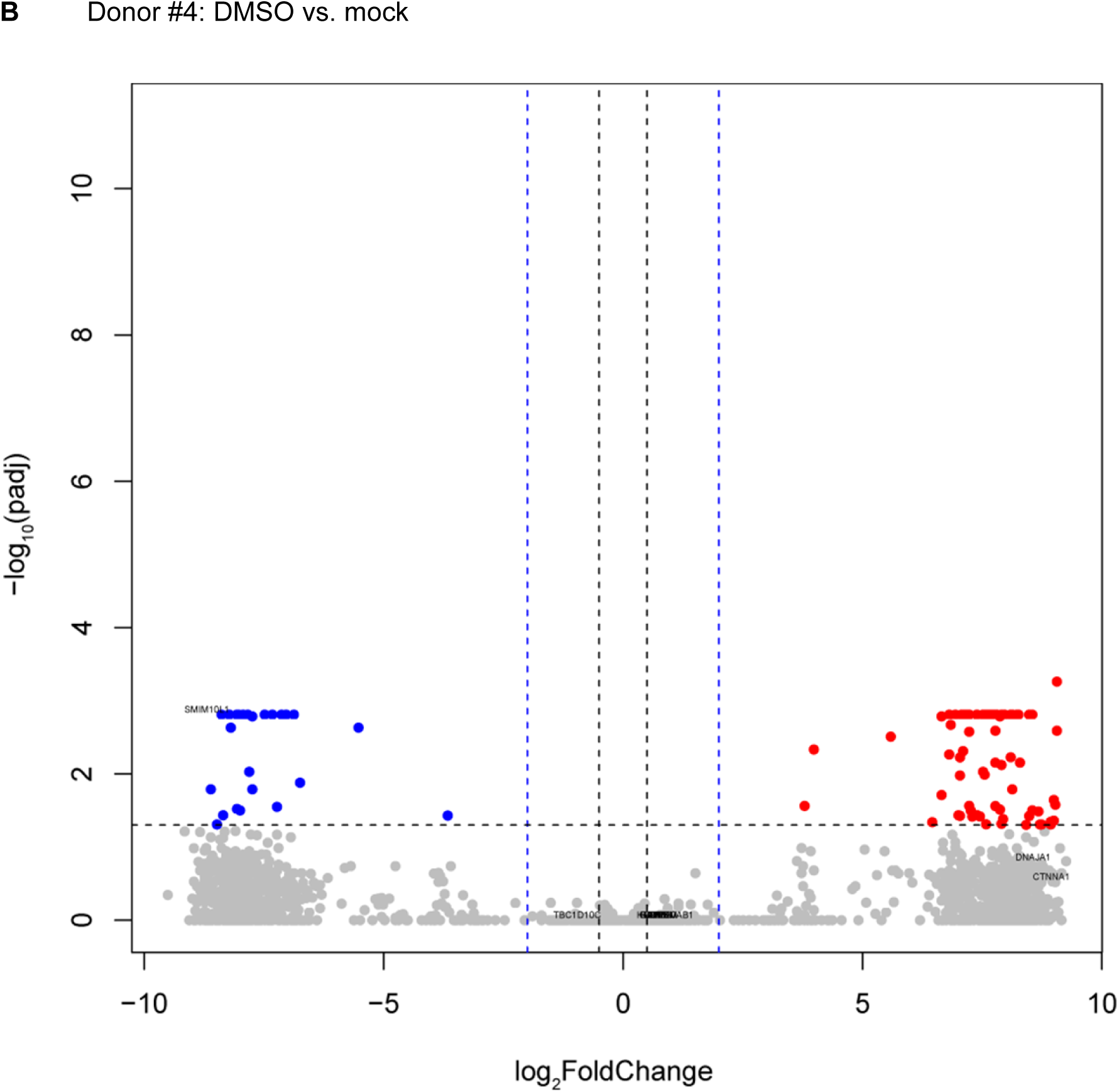
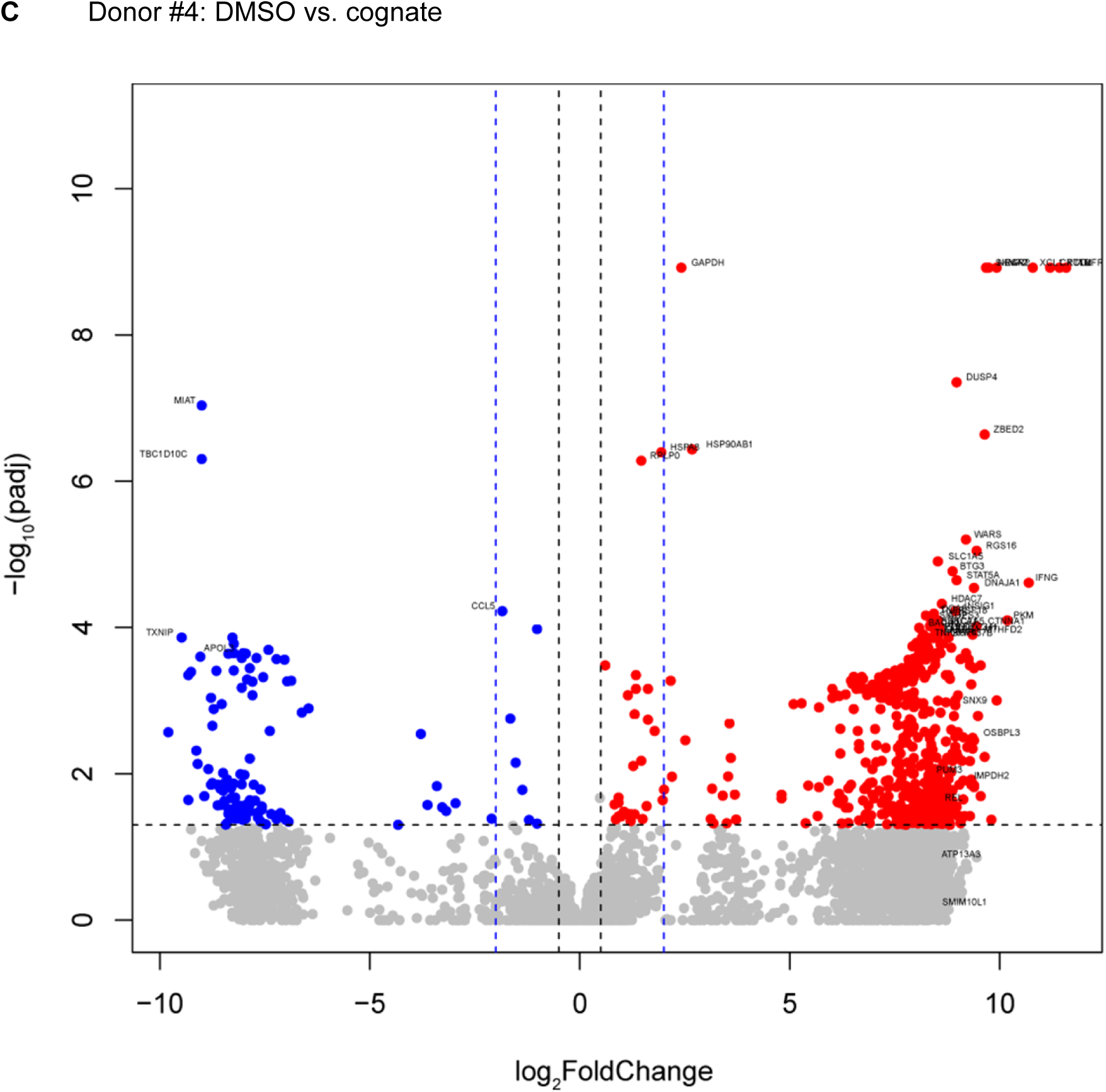
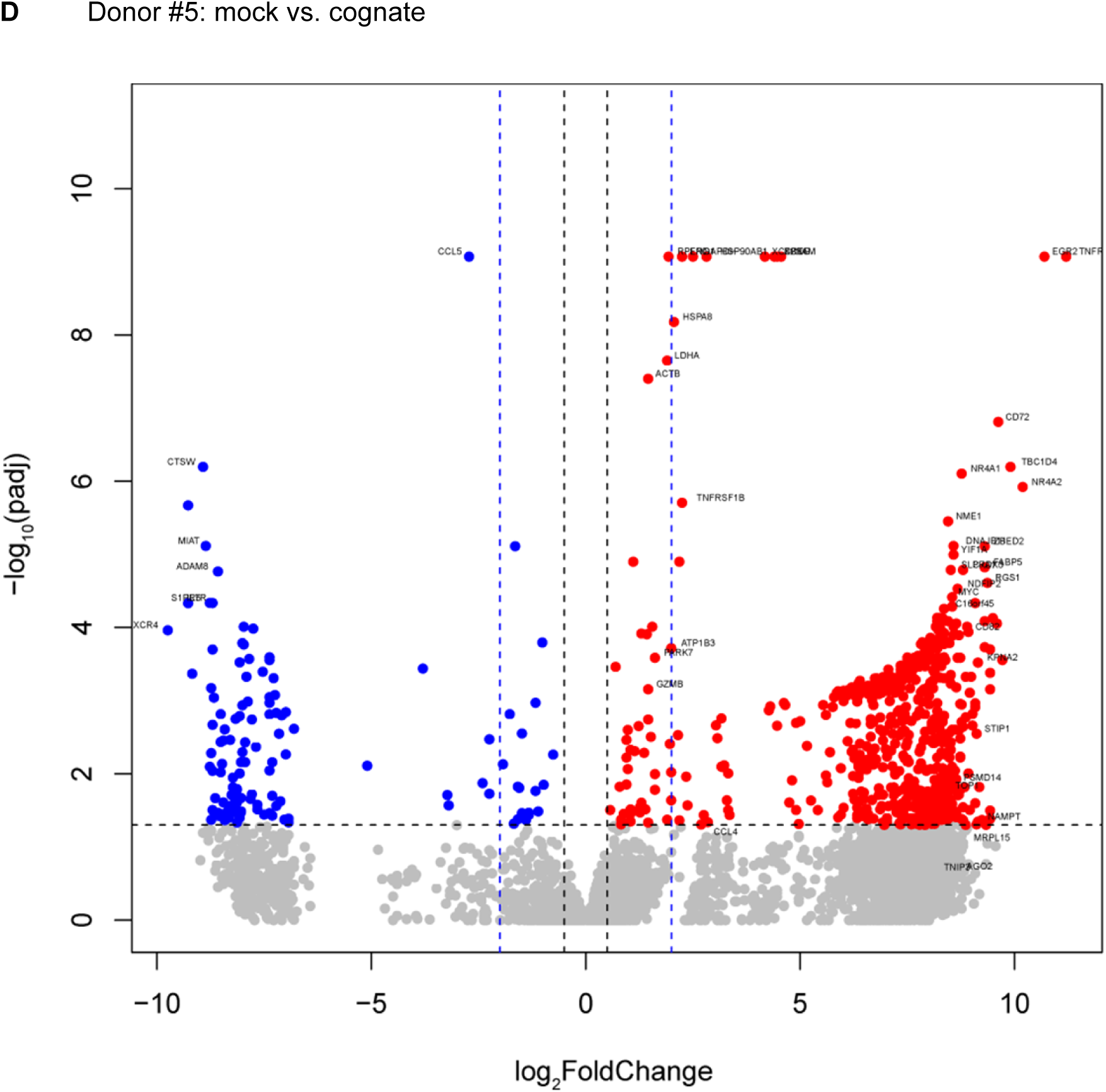
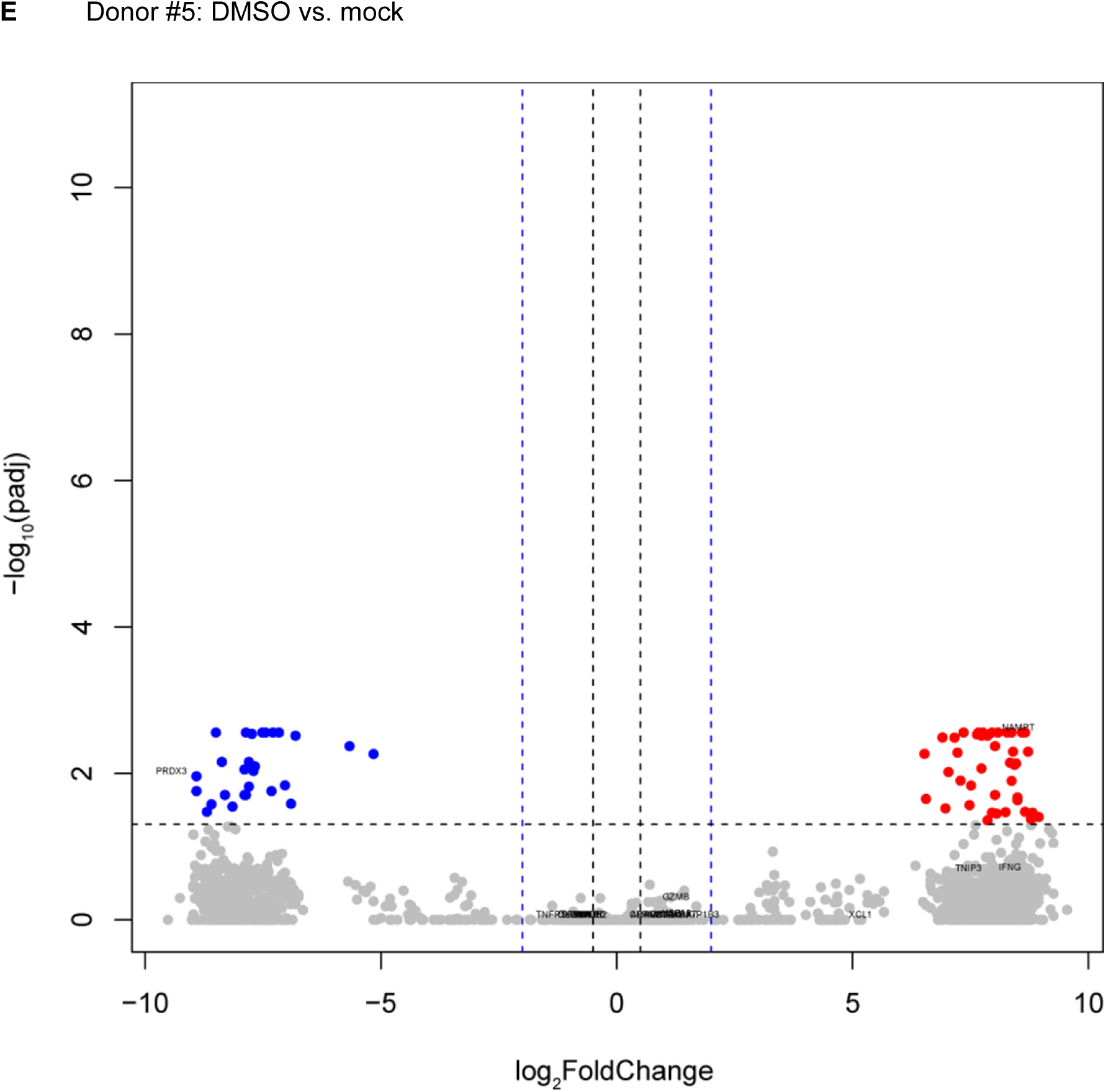
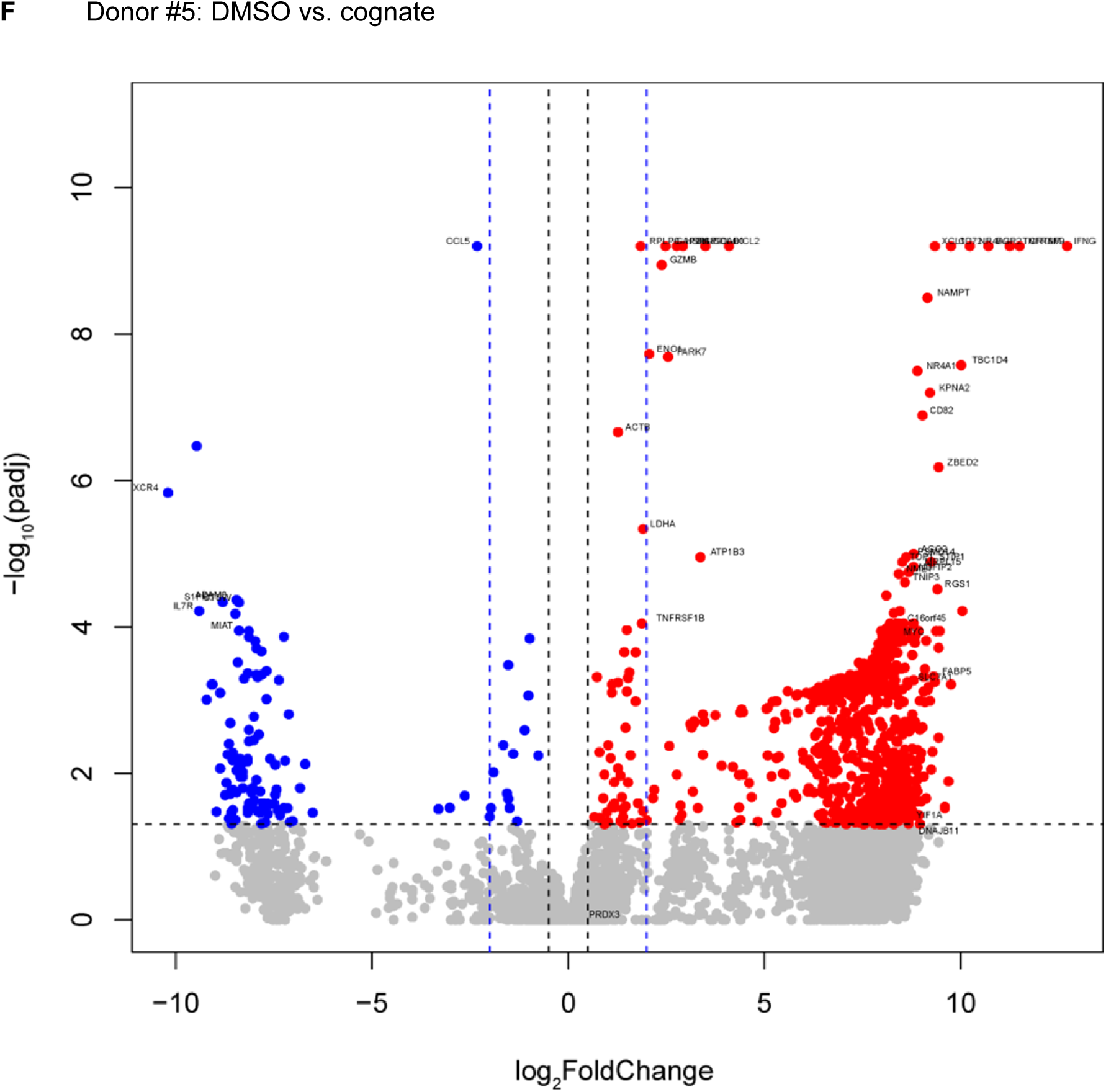
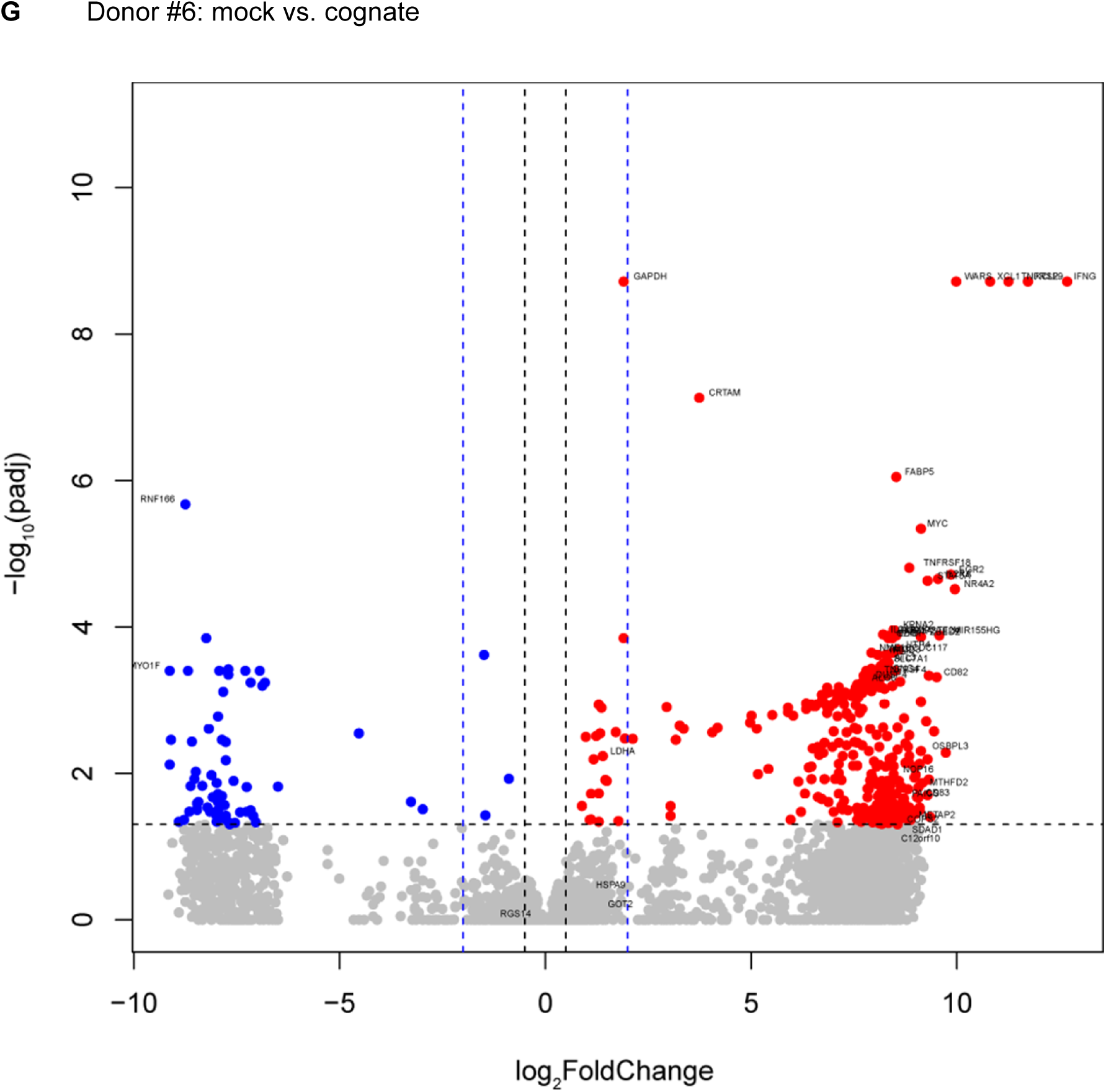
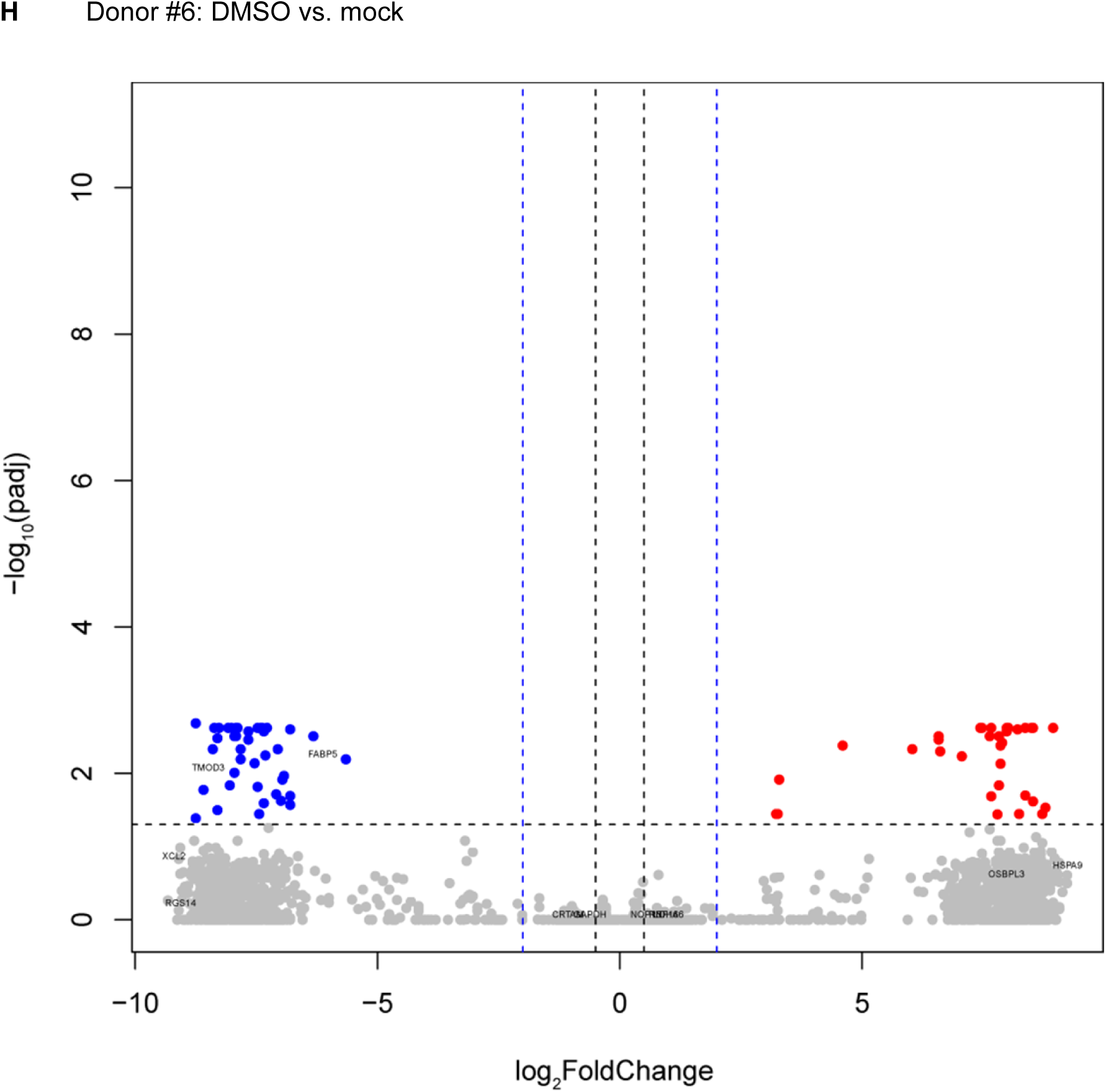
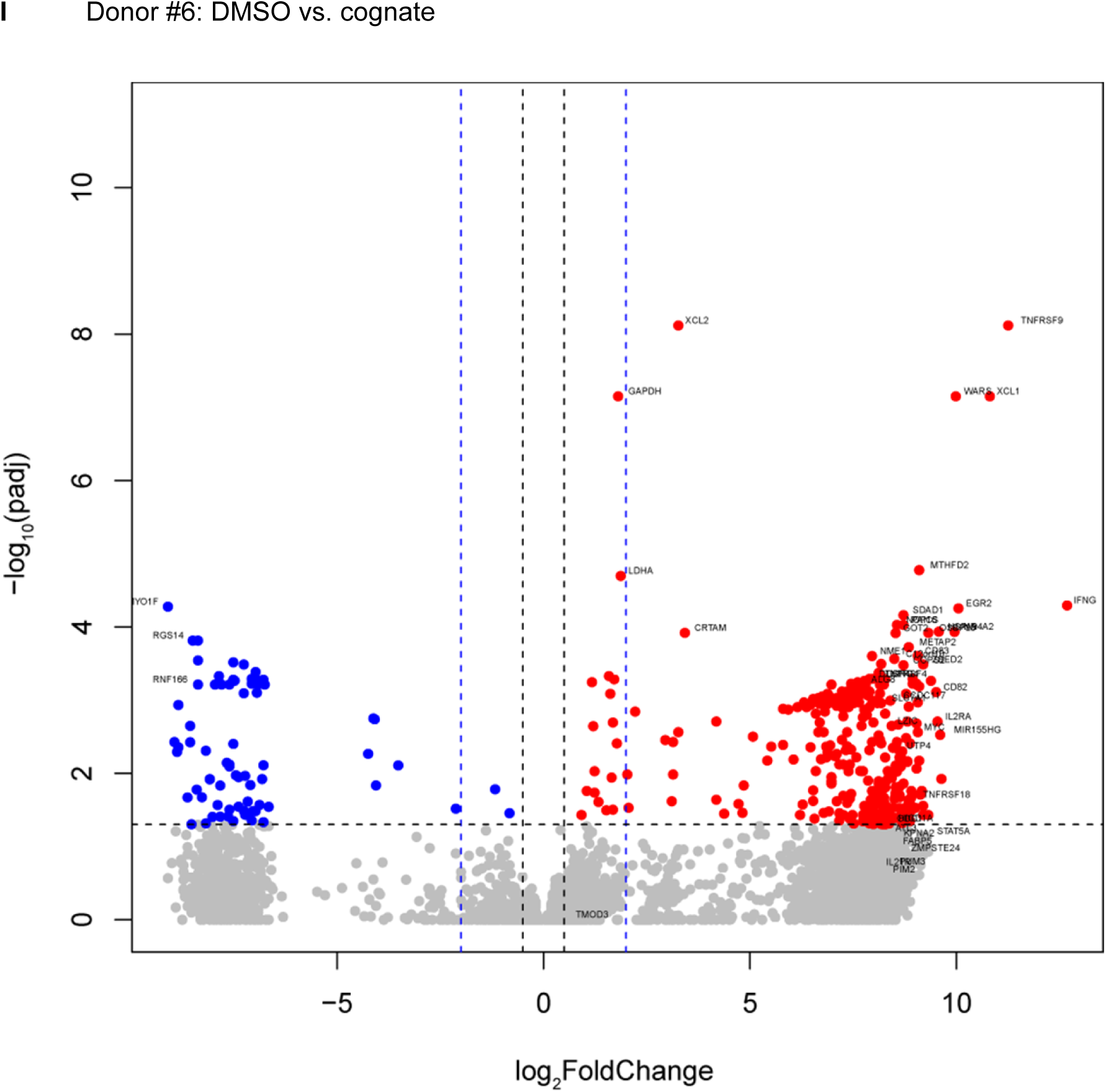
Comparison of gene expression between cognate peptide-stimulated and control-stimulated CMV pp65_495-503_-directed CD8^+^ T cells. (**A–I**) Volcano plots of scRNAseq data of CMV pp65_495-503_-directed cells from donors #4–6 are shown. Cells were single-cell-sorted from the dye-based CD8^+^ T cell activation assay following overnight incubation with peptide solvent (DMSO), mock peptide (Flu MP_58-66_), or cognate peptide (CMV pp65_495-503_) in the presence of autologous PBMCs for antigen presentation. The thresholds (fold change = 0.5 [black] and 4 [blue]; adjusted *p* = 0.05 [black]) are shown as dashed lines. Genes upregulated in cells responding to their cognate peptide are marked in red and downregulated genes in blue. The gene symbols for the combined top 50 ranked differentially expressed genes in both comparisons (DMSO vs. cognate and mock vs. cognate) are shown. Volcano plots comparing cells incubated with peptide solvent and mock peptide (DMSO vs. mock) are included as control conditions. Genes upregulated in cells stimulated with mock peptide are marked in red and downregulated genes in blue.

**Fig. S5:**
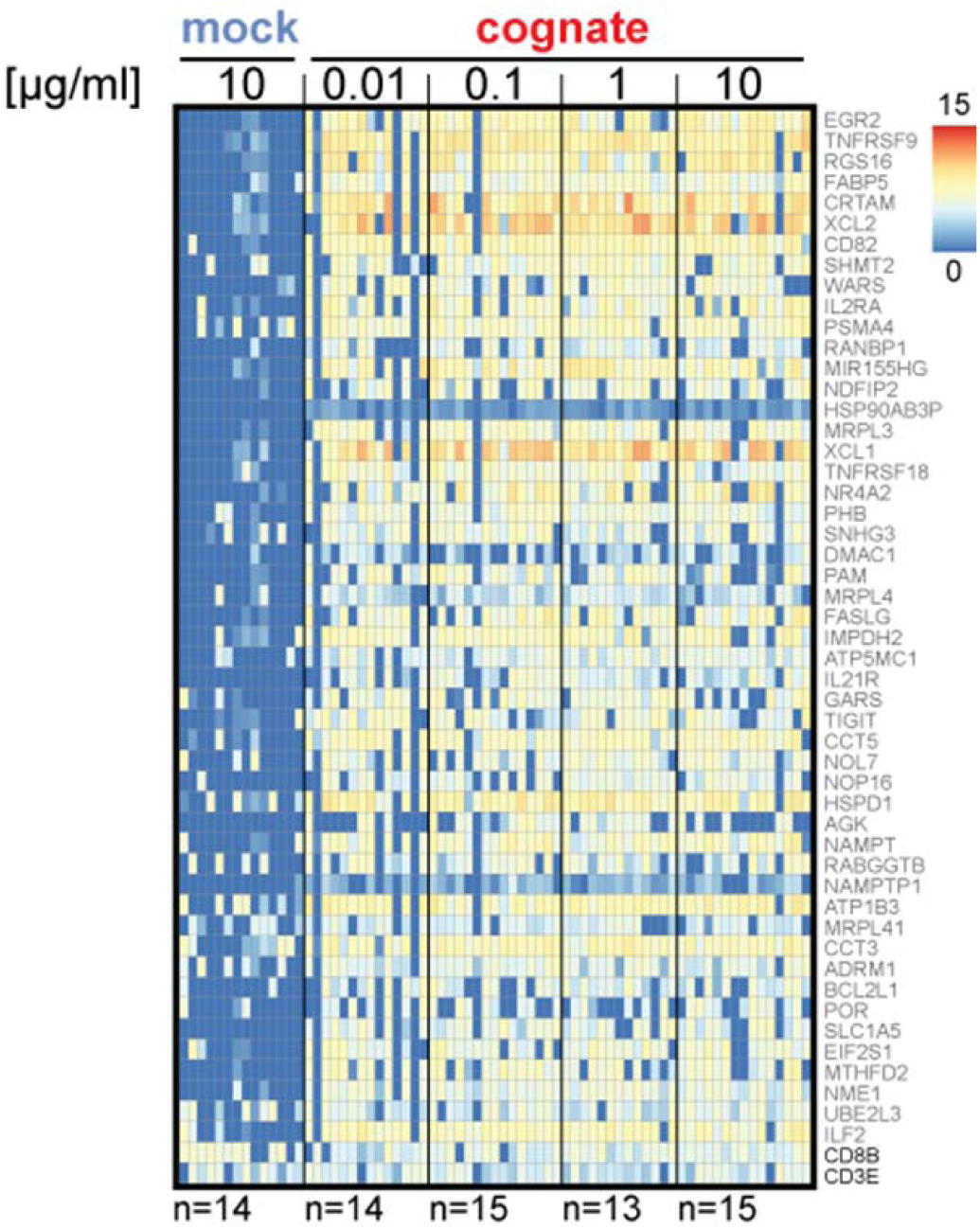
Discriminative ability of antigen-responsive cells across a wide range of cognate peptide concentrations. Heatmap showing gene expression in single-cell-sorted Flu MP_58-66_-directed cells incubated overnight with mock (IGRP_265-273_) or titrated amounts of cognate (Flu MP_58-66_) peptide in the dye-based CD8^+^ T cell activation assay using autologous PBMCs for antigen presentation. Results are shown for the top 50 ranked separator genes for cognate peptide-responsive CD8^+^ T cells of donor #1 and PBMC-based stimulation (see Table S5).

**Fig. S6:**
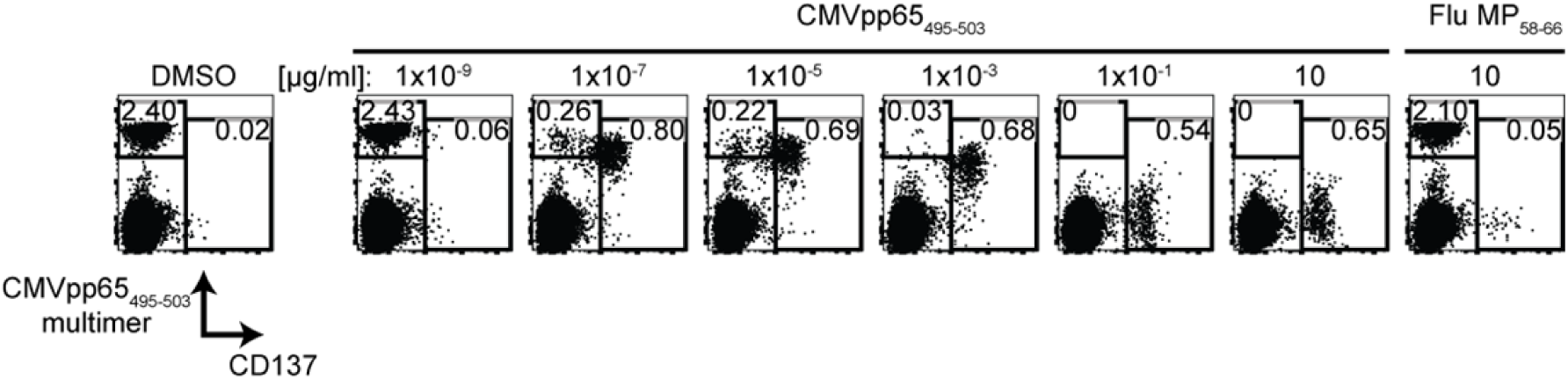
Impaired MHC class I multimer detection and upregulation of CD137 in CD8^+^ T cells responding to their cognate antigen. Representative flow cytometry dot plots of PBMCs stained with HLA-A2 multimers loaded with CMV pp65_495-503_ peptide after the PBMCs were incubated for 20 h in the presence of CMV pp65_495-503_ (cognate) or Flu MP_58-66_ peptide (mock) at the indicated concentrations. Plots show 0.5 × 10^4^ cells in the CD8 gate and the frequencies of multimer-stained cells (left gate) or CD137-expressing cells (right gate).

**Fig. S7:**
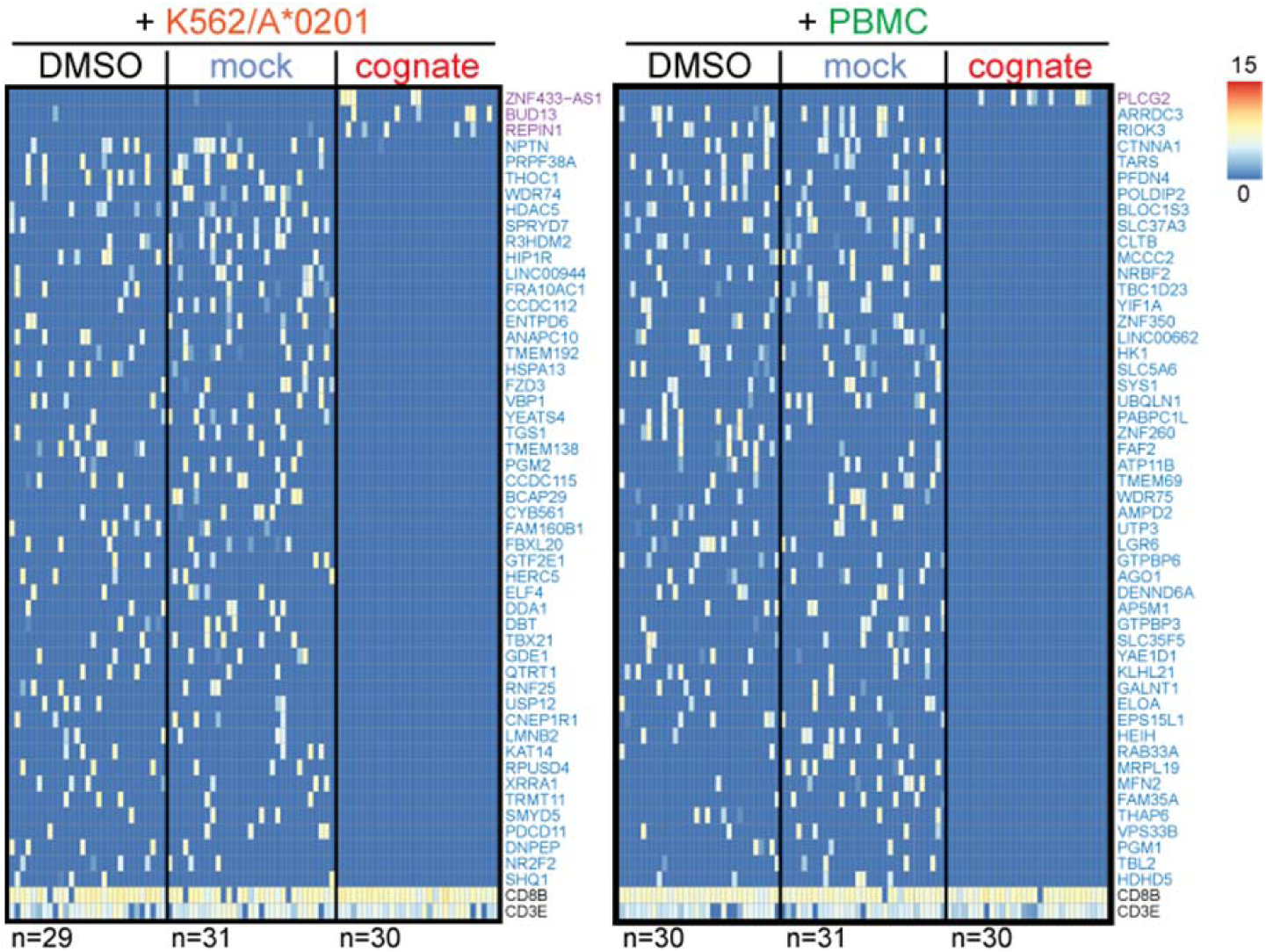
IGRP_265-273_-directed CD8^+^ T cells markedly differ from virus-directed cells in their response to cognate antigen. Heatmaps showing the top 50 ranked differentially expressed genes in autoantigen-directed CD8^+^ T cells incubated with cognate peptide (IGRP_265-273_) relative to control cells incubated with DMSO or mock peptide (Flu MP_58-66_). Cells were derived from a donor with type 1 diabetes and data were generated by scRNAseq. The top 50 genes were ranked separately for the K562/A*0201 (left) and autologous PBMC (right) antigen presentation conditions.

**Fig. S8:**
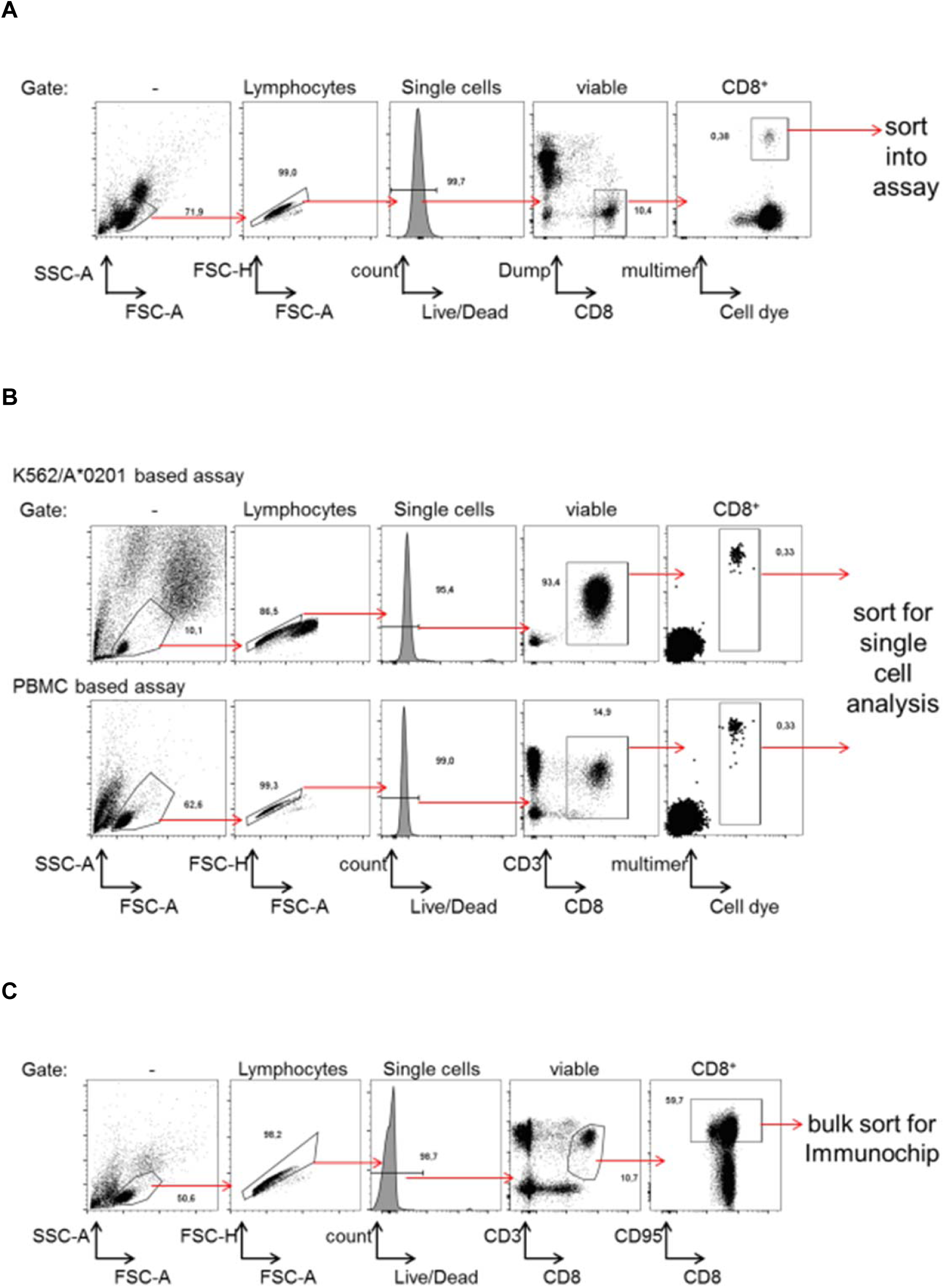
Gating strategies for the flow sorting experiments. Representative flow cytometry plots and gates are shown for sorting of (**A**) multimer-positive CD8^+^ T cells into the dye-based activation assay, (**B**) dye-stained antigen-directed CD8^+^ T cells from K562/A*0201- or PBMC-based assays for single-cell gene expression analysis, and (**C**) CD95^+^CD8^+^ memory T cells for immunochip analysis.

**Table S1:**
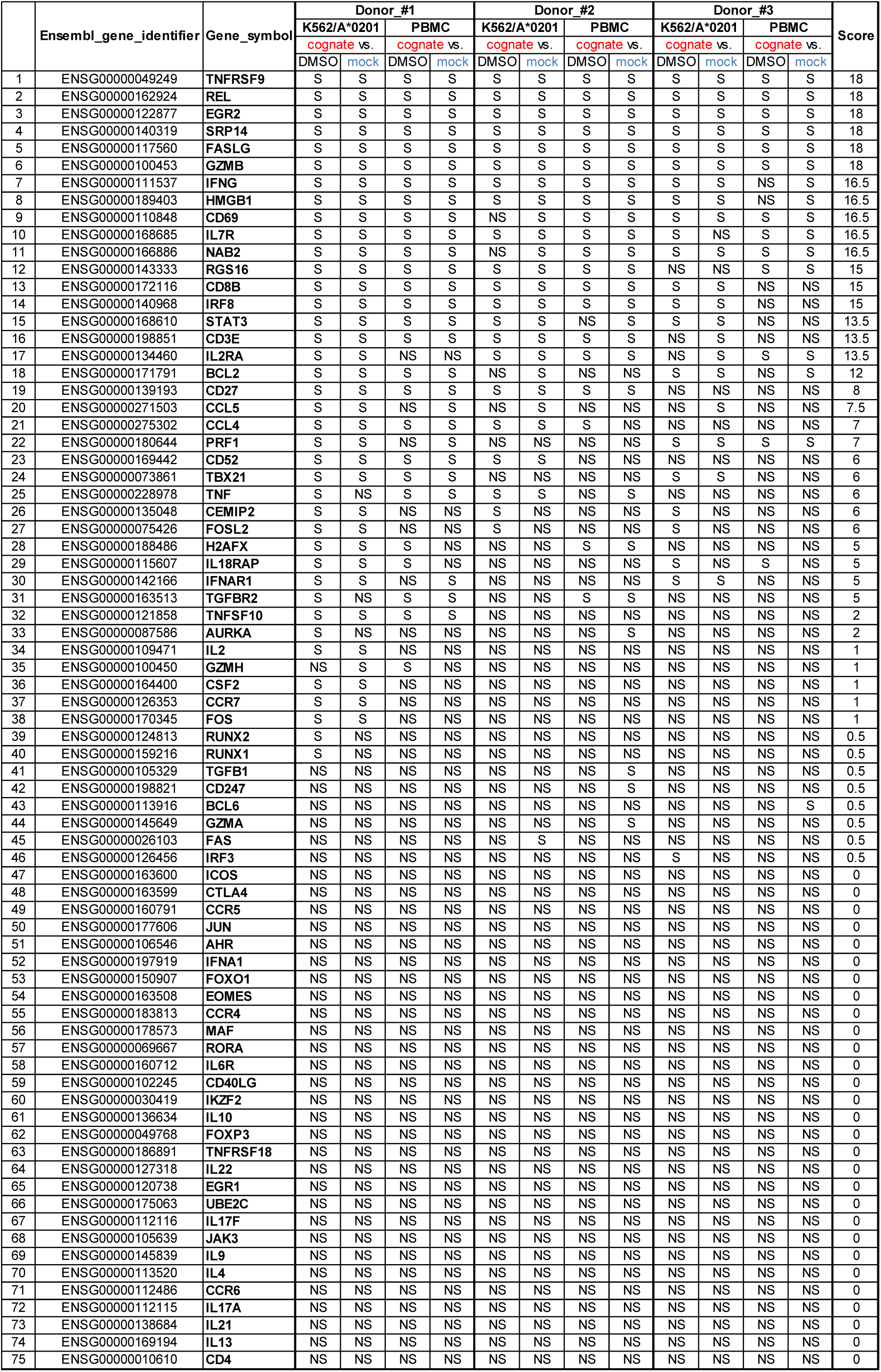
Overview of statistical results of the targeted gene expression analysis. qPCR-based single-cell gene expression data of Flu MP_58-66_-directed cells from donors #1–3 stimulated with cognate peptide or control stimuli (solvent DMSO or mock peptide) in the dye-based CD8^+^ T cell activation assay were compared using the Hurdle model (38). Genes were ranked using scores that considered the number of comparisons suggesting significantly different gene expression levels and the number of donors with statistically significant comparisons. Genes with the same score were ranked based on the delta for their median expression differences between the cognate and control stimuli. S = significant; NS = not significant.

**Table S2: T cell receptor information for the individual donors.**

See separate Excel file

**Table S3: Differentially expressed genes in cognate antigen-responsive CD8^+^ T cells from donor 1.**

See separate Excel file

**Table S4: Differentially expressed genes in cognate antigen-responsive CD8^+^ T cells from donors 4–6.**

See separate Excel file

**Table S5: Separator index and ranking of separator genes.**

See separate Excel file

**Table S6: Differentially expressed genes in CD137-expressing antigen-responsive cells.**

See separate Excel file

**Table S7: SVM-predicted Flu MP_58-66_-responsive CD8^+^ T cells.**

See separate Excel file

**Table S8: Differentially expressed genes in cognate antigen-stimulated CD8^+^ T cells from a donor with type 1 diabetes.**

See separate Excel file

**Table S9:**
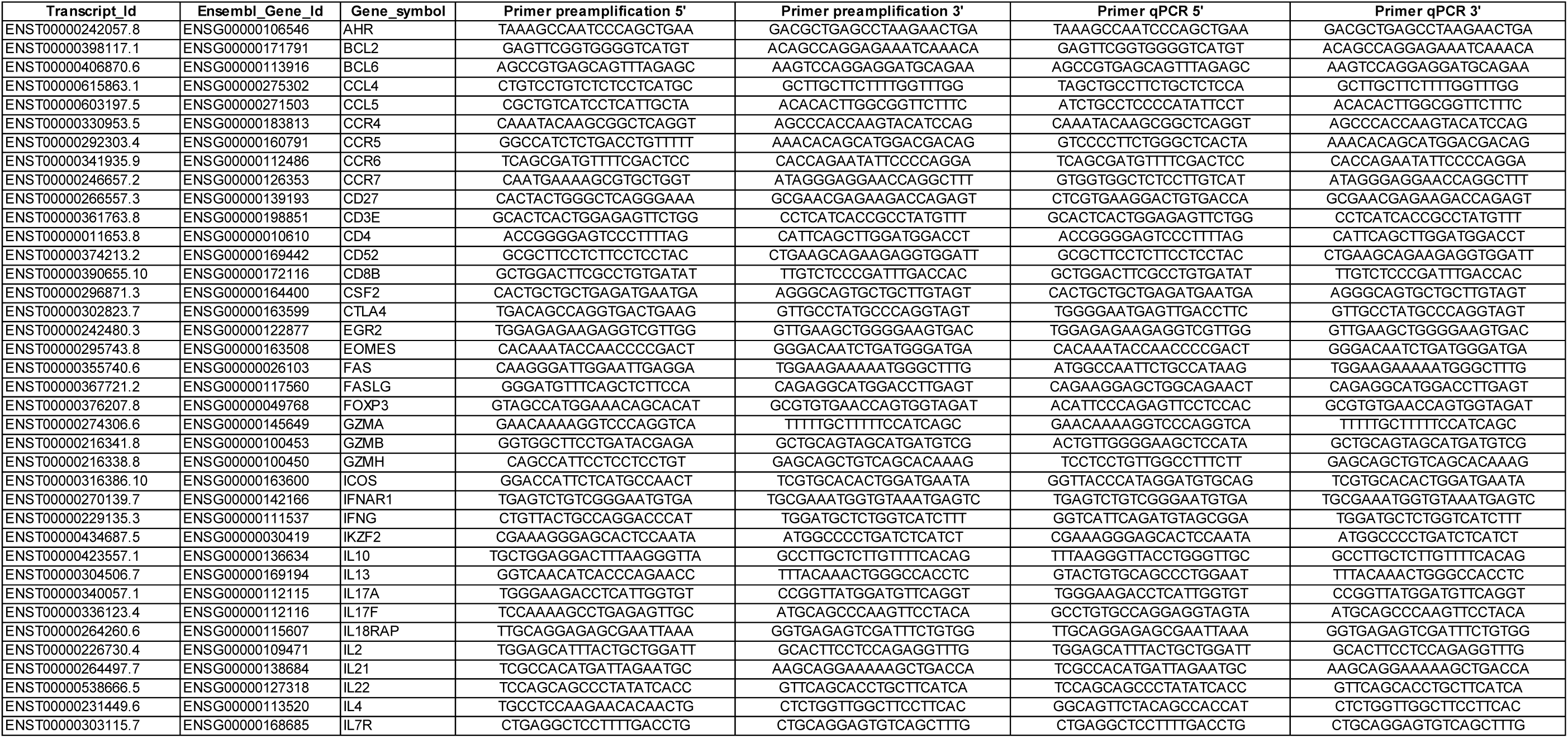
Primer pairs used for preamplification and qPCR.

## References

1. R. S. Andersen, P. Kvistborg, T. M. Frosig, N. W. Pedersen, R. Lyngaa, A. H. Bakker, C. J. Shu, P. Straten, T. N. Schumacher, S. R. Hadrup, Parallel detection of antigen-specific T cell responses by combinatorial encoding of MHC multimers. Nature protocols 7, 891–902 (2012).

2. W. W. Unger, J. Velthuis, J. R. Abreu, S. Laban, E. Quinten, M. G. Kester, S. Reker-Hadrup, A. H. Bakker, G. Duinkerken, A. Mulder, K. L. Franken, R. Hilbrands, B. Keymeulen, M. Peakman, F. Ossendorp, J. W. Drijfhout, T. N. Schumacher, B. O. Roep, Discovery of low-affinity preproinsulin epitopes and detection of autoreactive CD8 T-cells using combinatorial MHC multimers. Journal of autoimmunity 37, 151–159 (2011).

3. S. R. Hadrup, A. H. Bakker, C. J. Shu, R. S. Andersen, J. van Veluw, P. Hombrink, E. Castermans, P. Thor Straten, C. Blank, J. B. Haanen, M. H. Heemskerk, T. N. Schumacher, Parallel detection of antigen-specific T-cell responses by multidimensional encoding of MHC multimers. Nature methods 6, 520–526 (2009).

4. J. Glanville, H. Huang, A. Nau, O. Hatton, L. E. Wagar, F. Rubelt, X. Ji, A. Han, S. M. Krams, C. Pettus, N. Haas, C. S. L. Arlehamn, A. Sette, S. D. Boyd, T. J. Scriba, O. M. Martinez, M. M. Davis, Identifying specificity groups in the T cell receptor repertoire. Nature 547, 94–98 (2017).

5. P. Dash, A. J. Fiore-Gartland, T. Hertz, G. C. Wang, S. Sharma, A. Souquette, J. C. Crawford, E. B. Clemens, T. H. O. Nguyen, K. Kedzierska, N. L. La Gruta, P. Bradley, P. G. Thomas, Quantifiable predictive features define epitope-specific T cell receptor repertoires. Nature 547, 89–93 (2017).

6. H. C. Fan, G. K. Fu, S. P. Fodor, Expression profiling. Combinatorial labeling of single cells for gene expression cytometry. Science 347, 1258367 (2015).

7. G. Dolton, E. Zervoudi, C. Rius, A. Wall, H. L. Thomas, A. Fuller, L. Yeo, M. Legut, S. Wheeler, M. Attaf, D. M. Chudakov, E. Choy, M. Peakman, A. K. Sewell, Optimized Peptide-MHC Multimer Protocols for Detection and Isolation of Autoimmune T-Cells. Frontiers in immunology 9, 1378 (2018).

8. M. Wang, D. Windgassen, E. T. Papoutsakis, Comparative analysis of transcriptional profiling of CD3+, CD4+ and CD8+ T cells identifies novel immune response players in T-cell activation. BMC genomics 9, 225 (2008).

9. C. Kao, M. A. Daniels, S. C. Jameson, Loss of CD8 and TCR binding to Class I MHC ligands following T cell activation. International immunology 17, 1607–1617 (2005).

10. A. Alcover, B. Alarcon, V. Di Bartolo, Cell Biology of T Cell Receptor Expression and Regulation. Annual review of immunology 36, 103–125 (2018).

11. A. Viola, A. Lanzavecchia, T cell activation determined by T cell receptor number and tunable thresholds. Science 273, 104–106 (1996).

12. S. A. Valkenburg, T. M. Josephs, E. B. Clemens, E. J. Grant, T. H. Nguyen, G. C. Wang, D. A. Price, A. Miller, S. Y. Tong, P. G. Thomas, P. C. Doherty, J. Rossjohn, S. Gras, K. Kedzierska, Molecular basis for universal HLA-A*0201-restricted CD8+ T-cell immunity against influenza viruses. Proceedings of the National Academy of Sciences of the United States of America 113, 4440–4445 (2016).

13. Y. N. Naumov, E. N. Naumova, M. B. Yassai, K. Kota, R. M. Welsh, L. K. Selin, Multiple glycines in TCR alpha-chains determine clonally diverse nature of human T cell memory to influenza A virus. Journal of immunology 181, 7407–7419 (2008).

14. Y. F. Fuchs, G. W. Jainta, D. Kuhn, C. Wilhelm, M. Weigelt, A. Karasinsky, B. Upadhyaya, A. G. Ziegler, E. Bonifacio, Vagaries of the ELISpot assay: specific detection of antigen responsive cells requires purified CD8(+) T cells and MHC class I expressing antigen presenting cell lines. Clinical immunology 157, 216–225 (2015).

15. A. Brewitz, S. Eickhoff, S. Dahling, T. Quast, S. Bedoui, R. A. Kroczek, C. Kurts, N. Garbi, W. Barchet, M. Iannacone, F. Klauschen, W. Kolanus, T. Kaisho, M. Colonna, R. N. Germain, W. Kastenmuller, CD8(+) T Cells Orchestrate pDC-XCR1(+) Dendritic Cell Spatial and Functional Cooperativity to Optimize Priming. Immunity 46, 205–219 (2017).

16. A. Takeuchi, Y. Itoh, A. Takumi, C. Ishihara, N. Arase, T. Yokosuka, H. Koseki, S. Yamasaki, Y. Takai, J. Miyoshi, K. Ogasawara, T. Saito, CRTAM confers late-stage activation of CD8+ T cells to regulate retention within lymph node. Journal of immunology 183, 4220–4228 (2009).

17. Y. Pan, T. Tian, C. O. Park, S. Y. Lofftus, S. Mei, X. Liu, C. Luo, J. T. O’Malley, A. Gehad, J. E. Teague, S. J. Divito, R. Fuhlbrigge, P. Puigserver, J. G. Krueger, G. S. Hotamisligil, R. A. Clark, T. S. Kupper, Survival of tissue-resident memory T cells requires exogenous lipid uptake and metabolism. Nature 543, 252–256 (2017).

18. M. Wolfl, J. Kuball, W. Y. Ho, H. Nguyen, T. J. Manley, M. Bleakley, P. D. Greenberg, Activation-induced expression of CD137 permits detection, isolation, and expansion of the full repertoire of CD8+ T cells responding to antigen without requiring knowledge of epitope specificities. Blood 110, 201–210 (2007).

19. T. C. Wehler, M. Karg, E. Distler, A. Konur, M. Nonn, R. G. Meyer, C. Huber, U. F. Hartwig, W. Herr, Rapid identification and sorting of viable virus-reactive CD4(+) and CD8(+) T cells based on antigen-triggered CD137 expression. Journal of immunological methods 339, 23–37 (2008).

20. K. T. Coppieters, F. Dotta, N. Amirian, P. D. Campbell, T. W. Kay, M. A. Atkinson, B. O. Roep, M. G. von Herrath, Demonstration of islet-autoreactive CD8 T cells in insulitic lesions from recent onset and long-term type 1 diabetes patients. The Journal of experimental medicine 209, 51–60 (2012).

21. T. Takaki, M. P. Marron, C. E. Mathews, S. T. Guttmann, R. Bottino, M. Trucco, T. P. DiLorenzo, D. V. Serreze, HLA-A*0201-restricted T cells from humanized NOD mice recognize autoantigens of potential clinical relevance to type 1 diabetes. Journal of immunology 176, 3257–3265 (2006).

22. W. W. Unger, T. Pearson, J. R. Abreu, S. Laban, A. R. van der Slik, S. M. der Kracht, M. G. Kester, D. V. Serreze, L. D. Shultz, M. Griffioen, J. W. Drijfhout, D. L. Greiner, B. O. Roep, Islet-specific CTL cloned from a type 1 diabetes patient cause beta-cell destruction after engraftment into HLA-A2 transgenic NOD/scid/IL2RG null mice. PloS one 7, e49213 (2012).

23. Y. F. Fuchs, A. Eugster, S. Dietz, C. Sebelefsky, D. Kuhn, C. Wilhelm, A. Lindner, A. Gavrisan, J. Knoop, A. Dahl, A. G. Ziegler, E. Bonifacio, CD8(+) T cells specific for the islet autoantigen IGRP are restricted in their T cell receptor chain usage. Scientific reports 7, 44661 (2017).

24. S. Culina, A. I. Lalanne, G. Afonso, K. Cerosaletti, S. Pinto, G. Sebastiani, K. Kuranda, L. Nigi, A. Eugster, T. Osterbye, A. Maugein, J. E. McLaren, K. Ladell, E. Larger, J. P. Beressi, A. Lissina, V. Appay, H. W. Davidson, S. Buus, D. A. Price, M. Kuhn, E. Bonifacio, M. Battaglia, S. Caillat-Zucman, F. Dotta, R. Scharfmann, B. Kyewski, R. Mallone, Islet-reactive CD8(+) T cell frequencies in the pancreas, but not in blood, distinguish type 1 diabetic patients from healthy donors. Science immunology 3, (2018).

25. K. Matsuo, K. Kitahata, F. Kawabata, M. Kamei, Y. Hara, S. Takamura, N. Oiso, A. Kawada, O. Yoshie, T. Nakayama, A Highly Active Form of XCL1/Lymphotactin Functions as an Effective Adjuvant to Recruit Cross-Presenting Dendritic Cells for Induction of Effector and Memory CD8(+) T Cells. Frontiers in immunology 9, 2775 (2018).

26. B. G. Dorner, A. Scheffold, M. S. Rolph, M. B. Huser, S. H. Kaufmann, A. Radbruch, I. E. Flesch, R. A. Kroczek, MIP-1alpha, MIP-1beta, RANTES, and ATAC/lymphotactin function together with IFN-gamma as type 1 cytokines. Proceedings of the National Academy of Sciences of the United States of America 99, 6181–6186 (2002).

27. T. Miao, A. L. J. Symonds, R. Singh, J. D. Symonds, A. Ogbe, B. Omodho, B. Zhu, S. Li, P. Wang, Egr2 and 3 control adaptive immune responses by temporally uncoupling expansion from T cell differentiation. The Journal of experimental medicine 214, 1787–1808 (2017).

28. N. Du, H. Kwon, P. Li, E. E. West, J. Oh, W. Liao, Z. Yu, M. Ren, W. J. Leonard, EGR2 is critical for peripheral naive T-cell differentiation and the T-cell response to influenza. Proceedings of the National Academy of Sciences of the United States of America 111, 16484–16489 (2014).

29. J. C. Dudda, B. Salaun, Y. Ji, D. C. Palmer, G. C. Monnot, E. Merck, C. Boudousquie, D. T. Utzschneider, T. M. Escobar, R. Perret, S. A. Muljo, M. Hebeisen, N. Rufer, D. Zehn, A. Donda, N. P. Restifo, W. Held, L. Gattinoni, P. Romero, MicroRNA-155 is required for effector CD8+ T cell responses to virus infection and cancer. Immunity 38, 742–753 (2013).

30. E. Stelekati, Z. Chen, S. Manne, M. Kurachi, M. A. Ali, K. Lewy, Z. Cai, K. Nzingha, L. M. McLane, J. L. Hope, A. J. Fike, P. D. Katsikis, E. J. Wherry, Long-Term Persistence of Exhausted CD8 T Cells in Chronic Infection Is Regulated by MicroRNA-155. Cell reports 23, 2142–2156 (2018).

31. E. Lippert, D. L. Yowe, J. A. Gonzalo, J. P. Justice, J. M. Webster, E. R. Fedyk, M. Hodge, C. Miller, J. C. Gutierrez-Ramos, F. Borrego, A. Keane-Myers, K. M. Druey, Role of regulator of G protein signaling 16 in inflammation-induced T lymphocyte migration and activation. Journal of immunology 171, 1542–1555 (2003).

32. P. M. Gubser, G. R. Bantug, L. Razik, M. Fischer, S. Dimeloe, G. Hoenger, B. Durovic, A. Jauch, C. Hess, Rapid effector function of memory CD8+ T cells requires an immediate-early glycolytic switch. Nature immunology 14, 1064–1072 (2013).

33. R. Roge, J. Thorsen, C. Torring, A. Ozbay, B. K. Moller, J. Carstens, Commonly used reference genes are actively regulated in in vitro stimulated lymphocytes. Scandinavian journal of immunology 65, 202–209 (2007).

34. A. Grifoni, P. Costa-Ramos, J. Pham, Y. Tian, S. L. Rosales, G. Seumois, J. Sidney, A. D. de Silva, L. Premkumar, M. H. Collins, M. Stone, P. J. Norris, C. M. E. Romero, A. Durbin, M. J. Ricciardi, J. E. Ledgerwood, A. M. de Silva, M. Busch, B. Peters, P. Vijayanand, E. Harris, A. K. Falconar, E. Kallas, D. Weiskopf, A. Sette, Cutting Edge: Transcriptional Profiling Reveals Multifunctional and Cytotoxic Antiviral Responses of Zika Virus-Specific CD8(+) T Cells. Journal of immunology 201, 3487–3491 (2018).

35. C. M. Britten, R. G. Meyer, T. Kreer, I. Drexler, T. Wolfel, W. Herr, The use of HLA-A*0201-transfected K562 as standard antigen-presenting cells for CD8(+) T lymphocytes in IFN-gamma ELISPOT assays. Journal of immunological methods 259, 95–110 (2002).

36. S. Picelli, O. R. Faridani, A. K. Bjorklund, G. Winberg, S. Sagasser, R. Sandberg, Full-length RNA-seq from single cells using Smart-seq2. Nature protocols 9, 171–181 (2014).

37. E. Bonifacio, A. G. Ziegler, G. Klingensmith, E. Schober, P. J. Bingley, M. Rottenkolber, A. Theil, A. Eugster, R. Puff, C. Peplow, F. Buettner, K. Lange, J. Hasford, P. Achenbach, P. S. G. Pre, Effects of high-dose oral insulin on immune responses in children at high risk for type 1 diabetes: the Pre-POINT randomized clinical trial. Jama 313, 1541–1549 (2015).

38. A. McDavid, L. Dennis, P. Danaher, G. Finak, M. Krouse, A. Wang, P. Webster, J. Beechem, R. Gottardo, Modeling bi-modality improves characterization of cell cycle on gene expression in single cells. PLoS computational biology 10, e1003696 (2014).

39. P. V. Kharchenko, L. Silberstein, D. T. Scadden, Bayesian approach to single-cell differential expression analysis. Nature methods 11, 740–742 (2014).

40. M. I. Love, W. Huber, S. Anders, Moderated estimation of fold change and dispersion for RNA-seq data with DESeq2. Genome biology 15, 550 (2014).

41. A. Butler, P. Hoffman, P. Smibert, E. Papalexi, R. Satija, Integrating single-cell transcriptomic data across different conditions, technologies, and species. Nature biotechnology 36, 411–420 (2018).

42. M. J. T. Stubbington, T. Lonnberg, V. Proserpio, S. Clare, A. O. Speak, G. Dougan, S. A. Teichmann, T cell fate and clonality inference from single-cell transcriptomes. Nature methods 13, 329–332 (2016).

43. E. Alamyar, P. Duroux, M. P. Lefranc, V. Giudicelli, IMGT((R)) tools for the nucleotide analysis of immunoglobulin (IG) and T cell receptor (TR) V-(D)-J repertoires, polymorphisms, and IG mutations: IMGT/V-QUEST and IMGT/HighV-QUEST for NGS. Methods in molecular biology 882, 569–604 (2012).

